# Phosphate Deficiency Attenuates SnRK1 Activity to Modulate the Phosphate Starvation Response in *Arabidopsis thaliana*

**DOI:** 10.64898/2026.10.09.758112

**Authors:** Antonio Corbalán-Acedo, Adela Villasante-Fernández, Anthony Artins, Ana Confraria, Liliana Ferreira, Cecilia Costigliolo-Rojas, Antoni Garcia-Molina, Elena Baena-González, Alisdair R Fernie, Alejandro Ferrando, Borja Belda-Palazón

## Abstract

- Phosphate starvation requires extensive metabolic and developmental reprogramming, yet how this response is coordinated with cellular energy status remains poorly understood. The energy sensor SnRK1 regulates plant growth and metabolism in response to energy availability, but its contribution to the regulation and deployment of the phosphate starvation response (PSR) remains unclear.
- To address this, we combined physiological, proteomic and metabolomic analyses with genetic manipulation of SnRK1 activity *in Arabidopsis thaliana* to determine how SnRK1 influences plant responses to phosphate starvation.
- We found that phosphate starvation reduced SnRK1 activity and activation-loop (T-loop) phosphorylation, while altering SnRK1 activity changed the timing and magnitude of physiological, proteomic and metabolic responses to phosphate limitation. Reduced SnRK1 activity was generally associated with earlier and stronger deployment of PSR-associated responses, whereas increased SnRK1 activity delayed or attenuated several components of the response. These effects were tissue- and time-dependent and were accompanied by distinct metabolic configurations.
- Our findings support a model in which SnRK1 acts as a quantitative metabolic brake on the PSR, coordinating the metabolic investment required for phosphate-starvation adaptation with growth.

## Introduction

Phosphorus is an essential macronutrient required for virtually all aspects of plant growth and metabolism. As inorganic phosphate (Pi), it is a structural component of nucleic acids and phospholipids and is central to energy metabolism through its incorporation into ATP and other phosphorylated metabolites. However, despite its abundance in many soils, Pi is often poorly available to plants because of its low mobility and its tendency to form insoluble complexes with metal ions (Raghothama, 1999; Vance *et al*., 2003; Cooper *et al*., 2011; Lambers & Plaxton, 2015; Hallama *et al*., 2018). Consequently, plants frequently experience Pi limitation and have evolved a complex set of physiological, developmental and metabolic adaptations collectively known as the phosphate starvation response (PSR) (Dissanayaka et al., 2021; Paz-Ares et al., 2022).

The PSR involves coordinated changes aimed at increasing Pi acquisition, mobilization and recycling while adjusting plant growth and metabolism to limited Pi availability. These responses include the induction of high-affinity Pi transporters and phosphatases, remodeling of membrane lipids to reduce cellular Pi demand, changes in root system architecture, and extensive reprogramming of primary and secondary metabolism (Srivastava *et al*., 2018; Roychowdhury *et al*., 2023). The MYB-CC transcription factor PHOSPHATE STARVATION RESPONSE1 (PHR1) acts as a central regulator of this program, controlling a large fraction of the transcriptional response to Pi limitation and coordinating diverse processes involved in Pi homeostasis and adaptation (Nilsson *et al*., 2007; Bustos *et al*., 2010; Mangalakkadan *et al*., 2026). PHR1 activity is itself tightly regulated by Pi-dependent signalling mechanisms, including interactions with SPX-domain proteins and post-translational modifications. Importantly, however, the PSR is not simply a passive response to nutrient scarcity. Successful adaptation to Pi limitation requires extensive transcriptional, metabolic and developmental reprogramming, collectively demanding a substantial investment of carbon, ATP and reducing power. From an energetic perspective, therefore, the PSR cannot be regarded simply as a starvation response, but rather as an active adaptive program with a significant metabolic cost. How this substantial metabolic investment is coordinated with cellular energy status remains an important question (Dissanayaka et al., 2021; Roychowdhury et al., 2023).

A central regulator of cellular energy homeostasis in plants is SUCROSE NON-FERMENTING1-RELATED PROTEIN KINASE1 (SnRK1), the plant orthologue of AMPK in animals and SNF1 in yeast. SnRK1 is a conserved heterotrimeric protein kinase complex whose activity in *Arabidopsis thaliana* is primarily mediated by the catalytic α-subunits SnRK1α1 and SnRK1α2, in association with regulatory β and βγ subunits (Baena-González *et al*., 2007; Margalha *et al*., 2016). Under conditions associated with energy limitation or stress, SnRK1 promotes extensive transcriptional and metabolic reprogramming aimed at restoring energy balance by activating catabolic and ATP-producing processes while restricting biosynthetic activity and growth. SnRK1 therefore coordinates cellular energy status with metabolic investment, limiting energy-demanding processes when resources are scarce. Nevertheless, its role cannot simply be viewed as growth suppressive, as SnRK1 activity is also required for appropriate nutrient redistribution and development, and severe depletion of SnRK1 function is detrimental to plant growth (Jamsheer K *et al*., 2021).

This central role in coordinating energy status with metabolic investment creates an apparent paradox during phosphate starvation. Pi limitation compromises fundamental metabolic processes and may impose energetic constraints, conditions that might be expected to favour SnRK1 activation (Peixoto & Baena-González, 2022). At the same time, the establishment of the phosphate starvation response (PSR) requires substantial metabolic and developmental investment (Dissanayaka et al., 2021; Roychowdhury et al., 2023). A similarly complex relationship has been reported between Pi availability and TOR signalling. Studies have provided evidence for TOR inhibition during the early root response to Pi deficiency, whereas subsequent work has implicated TOR activity in promoting PHR1 stability and PSR-associated transcriptional reprogramming under prolonged Pi limitation (Cho *et al*., 2023; Awasthi *et al*., 2024). Together, these findings suggest that the relationship between central metabolic signalling pathways and the PSR cannot be explained by a simple nutrient-starvation paradigm.

Previous studies have reported complex and apparently divergent changes in SnRK1 during Pi starvation. Fragoso *et al*. reported an overall decrease in SnRK1 activity in Pi-starved leaves, accompanied by differential regulation of the two major catalytic subunits, including a marked reduction in SnRK1α2 protein abundance and an apparent increase in SnRK1α1 activity, as assessed by *in vitro* kinase assays using protein extracts from SnRK1α1-overexpressing plants (Fragoso *et al*., 2009). Subsequent work identified high-molecular-weight fractions with increased SnRK1 activity during Pi starvation, in which SnRK1α1 was the predominant catalytic subunit, together with several regulatory SnRK1 subunits (Trejo-Fregoso *et al*., 2022). The same study further showed that SnRK1 can phosphorylate PHR1 at Ser11, negatively affecting its transcriptional activity. These findings indicate that Pi starvation does not simply switch SnRK1 signalling on or off but rather induces a complex remodelling of SnRK1 activity and composition. However, these studies primarily focused on the regulation and composition of SnRK1 complexes or on the identification of potential molecular targets. Whether this remodelling of SnRK1 activity functionally contributes to whole-plant adaptation to Pi starvation, including the establishment and execution of the PSR, remains largely unexplored.

Here, we show that phosphate starvation is associated with a consistent attenuation of canonical SnRK1 activity *in vivo*. Using complementary genetic approaches, we further show that reduced SnRK1 activity promotes, whereas elevated SnRK1 activity restrains, key physiological responses to Pi limitation. Proteomic and metabolomic analyses reveal that SnRK1 influences the timing and magnitude of the response to phosphate starvation and shapes tissue-specific metabolic adaptation. Together, our findings identify attenuation of SnRK1 signalling as an important component of the adaptive response to Pi starvation and suggest that reduced SnRK1 activity facilitates the metabolic and developmental reprogramming required for PSR establishment.

## Materials and methods

### Plant materials and growth conditions

*Arabidopsis thaliana* ecotype Columbia-0 (Col-0) was used as the wild-type control. All mutant and transgenic lines used in this study are listed in Table **S1**. Unless otherwise indicated, plants were grown under long-day conditions (16 h light/8 h dark; 22°C/18°C) with a light intensity of 100-120 μmol m⁻² s⁻¹. Surface-sterilized seeds were sown on half-strength Murashige and Skoog (½ MS) medium with vitamins and MES (Duchefa M0255) supplemented with 1.03 g/L of MES and 0.8% (w/v) Bacto^TM^ Agar (BD) or Noble Agar, Ultrapure (Thermo Scientific) (pH 5.7). For phosphate-deficient (-Pi) conditions, the same medium was prepared using phosphate-free ½ MS Basal Salts (Modification 11; Plant Cell Labs), supplemented with 1.28 g/L of MES. Where indicated, seedlings were incubated for the specified period in phosphate-free liquid ½ MS medium. Plates were sealed with Micropore™ tape, stratified in darkness at 4°C for 2 days, and then transferred to the growth chamber.

### Cloning procedures and plant transformation

To generate SnRK1α1/KIN10- and SnRK1α2/KIN11-targeting artificial microRNA constructs, we designed artificial microRNAs using the Web MicroRNA Designer available at http://wmd3.weigelworld.org/cgi-bin/webapp.cgi?page=Home;project=stdwmd (Ossowski *et al*., 2008). We then mutagenised a pHBT95-pri319 construct (pri319 - primary miR319a) (Martinho *et al*., 2015) to generate an artificial microRNA targeting SnRK1α1/KIN10 and SnRK1α2/KIN11 (amiRKIN10/11), following the described protocol (Schwab *et al*., 2006). After confirmation by Sanger sequencing, the pri319 sequence carrying the newly generated amiRKIN10/11 was then excised by restriction digestion with BamHI and PstI and ligated into a pCB302 backbone (Xiang *et al*., 1999) using T4 DNA ligase according to the manufacturer’s instructions.

To generate the SnRK1α2-OE constructs, we excised the SnRK1α2/KIN11 coding sequence from a pHBT95-KIN11 construct (Baena-González *et al*., 2007) by restriction digestion with BamHI and StuI and ligated it into pCB302, as described above.

In both cases, constructs were transformed into Agrobacterium tumefaciens GV3101 and positive colonies were selected by antibiotic resistance (kanamycin). *Agrobacterium tumefaciens* carrying either the pCB302-amiRKIN10/11 or the pCB302-KIN11 construct was used to transform wildtype *A. thaliana* Col-0 plants by flower dipping (Clough & Bent, 1998). We selected independent lines with T2 antibiotic resistance of ∼75% and then propagated the lines until homozygosity.

### Primary root growth assays

For primary root growth assays, surface-sterilized seeds were germinated on ½ MS medium and grown for 5 days under long-day conditions, except for *sesquiα2-1* seedlings which were grown for 6 days on ½ MS supplemented with Basta (25 µg/mL) to allow comparable initial seedling development. Seedlings were then transferred to fresh ½ MS plates supplemented with or without Pi and grown for an additional 6 days. At the end of the treatment, plates were photographed and primary root growth was quantified from the scanned images using Fiji/ImageJ (Schindelin *et al*., 2012).

### Anthocyanin and soluble Pi content analyses

Anthocyanin and soluble Pi content were analysed as described for the root growth assays, following 4 to 7 days of treatment with or without Pi. Anthocyanin content was determined as described previously (Nakata & Ohme-Takagi, 2014). Soluble Pi content was determined using the ascorbate-molybdate colorimetric method described by Ames (1966), with minor modifications. Briefly, tissue was extracted in 1% acetic acid, and extracts were reacted with freshly prepared ascorbate-molybdate reagent at 37°C for 40 min. Absorbance was measured at 820 nm, and Pi content was calculated from an orthophosphate standard curve.

### SnRK1 activity reporter analysis

SnRK1 activity was monitored *in vivo* using the genetically encoded ASP-SPARK reporter as described previously (Safi *et al*., 2023). Seedlings were grown for 6 days under Pi-sufficient (control) or Pi-deficient (-Pi) conditions, and root tips and leaves were imaged separately. Confocal imaging was performed using a Stellaris 8 laser scanning confocal microscope (Leica Microsystems) equipped with a HC PL APO 40×/1.30 OIL objective and HyD detectors. eGFP fluorescence was excited at 490 nm and detected between 500 and 530 nm, whereas mCherry fluorescence was excited at 586 nm and detected between 600 and 630 nm. For multicolour imaging, fluorophores were acquired sequentially to avoid spectral overlap. Pinhole size was adjusted to 1 Airy unit for each channel. Images were acquired using Leica LAS X software and processed using Fiji/ImageJ. SnRK1 activity was quantified by calculating the SPARK ratio as described by Safi *et al*., (2023).

### Protein extraction and immunoblot analysis

Approximately 100 mg of frozen *A. thaliana* tissue was ground in liquid nitrogen and homogenized in two volumes of 2× Laemmli extraction buffer (126 mM Tris-HCl, pH 6.8, 4% SDS, 20% glycerol, 0.05% bromophenol blue and 4% β-mercaptoethanol). Samples were incubated on ice for 20-30 min, boiled at 95°C for 10-15 min and centrifuged at 12,000 × *g* for 15 min. Supernatants were used as total protein extracts. Equal volumes were separated by SDS-PAGE and transferred to PVDF membranes (Cytiva). Membranes were blocked with 5% (w/v) non-fat dry milk in 1× TBS-T (Tris-Buffered Saline with 0.05% Tween-20) and incubated overnight at 4°C with primary antibodies, followed by HRP-conjugated secondary antibodies. Signals were detected by quimioluminescence and quantified using Fiji/ImageJ. Antibodies are listed in Table **S3**.

### RNA extraction and RT-qPCR analysis

For gene expression analyses, total RNA was extracted from whole *A. thaliana* seedlings using the NZY Total RNA Isolation kit (NZYtech) according to the manufacturer’s instructions, treated with TURBO DNA-free™ Kit (Invitrogen, Thermo Fisher Scientific), and subsequently purified by 6 M LiCl precipitation. RNA concentration and purity were assessed using a NanoDrop™ One spectrophotometer (Thermo Fisher Scientific), and first-strand cDNA was synthesized from 1 μg total RNA using the PrimeScript™ RT reagent Kit (Takara Bio Inc.) according to the manufacturer’s instructions. RT-qPCR was performed using HOT FIREPol® EvaGreen® qPCR Mix Plus (Solis Biodyne) in 10 μL reactions containing 0.4 μL cDNA and 500 nM of each primer on a QuantStudio™ 3 Real-Time PCR System (Applied Biosystems). Relative transcript abundance was calculated using the 2^-ΔCt^ method, with *PDF2* as the reference gene. Each biological replicate was analysed in triplicate. Primer sequences are listed in Table **S2**.

### Proteomic and metabolomic analyses

For proteomic and metabolomic profiling, seedlings were initially grown under standard conditions for 10 days and subsequently transferred to fresh ½ MS plates containing either sufficient (control) or no phosphate (-Pi). Shoots and roots were harvested separately after 2 or 4 days of treatment and processed for proteomic and metabolomic analyses. For proteomic analysis, approximately 100 mg of frozen tissue were extracted in RIPA buffer (Thermo Scientific) supplemented with 50 µM MG132 (MedChemexpress), phosphatase inhibitor cocktails 2 and 3 (Sigma-Aldrich), and Complete Mini EDTA-free protease inhibitor (Roche). Extracts were heated for 15 min in Laemmli buffer and stored at -80°C until analysis. 10 µg of protein were processed using an SP3-based digestion protocol with minor modifications (Moggridge *et al*., 2018; Hughes *et al*., 2019), reduced with DTT, alkylated with iodoacetamide and digested overnight with trypsin. Peptides were analysed by LC-MS/MS on an Evosep One coupled to a timsTOF fleX mass spectrometer (Bruker) using a 30 SPD chromatographic method and diaPASEF acquisition. Raw data were processed using DIA-NN 1.8 against an *Arabidopsis thaliana* sequence database (Demichev *et al*., 2019). Protein abundance matrices generated by DIA-NN were subsequently analysed using MSstatsShiny for differential protein abundance analysis (Kohler *et al*., 2023). Unique peptides were used for protein quantification, and a Q-value cutoff of 0.01 was applied. Median equalization was used for normalization, and the top 36 features were selected for protein summarization using the Tukey’s median polish (TMP) method. Missing values were treated as censored and model-based imputation was applied. Data processing, statistical analyses and visualization were performed in R v4.6.0 (R Core Team, 2026).

Metabolites were extracted from 50 mg of frozen, and finely ground shoot or root material using 1 mL of pre-cooled MTBE (methyl-tert-butyl ether) extraction buffer. After extraction, phase separation was performed by adding 0.5 mL of water (3:1, v/v) containing 0.2 mg/mL ribitol as an internal standard, following the procedure described by Giavalisco et al., (2011). The polar phase containing primary metabolites was collected, concentrated, and derivatized using *N*-methyl-*N*-trimethylsilyltrifluoroacetamide (MSTFA). The derivatized samples were subsequently analysed by gas chromatography coupled to time-of-flight mass spectrometry (GC-TOF-MS) using an Agilent 7890N gas chromatograph coupled to a Pegasus HT TOF mass spectrometer (LECO), as described by (Lisec *et al*., 2006). Peak detection, retention-time alignment, and metabolite identification based on comparison with the available spectral library were performed using the TargetSearch R package (Cuadros-Inostroza *et al*., 2009). Metabolite abundance was estimated from the corresponding peak intensities and normalized sequentially to the signal detected in extraction blanks, the ribitol internal standard, and the fresh weight (FW) of the sample.

### General statistical analyses

Statistical analyses were performed using GraphPad Prism 8 (GraphPad Software). Data are presented as mean ± SEM. Statistical significance was assessed by one-way or two-way ANOVA followed by Tukeýs or Dunnett’s multiple comparison tests, or by Student’s *t*-test when appropriate. Details of the statistical tests, sample sizes and significance levels are provided in the corresponding figure legends.

## Results

### Phosphate starvation is associated with attenuation of SnRK1 activity *in vivo*

Given the complex changes in SnRK1 activity and composition previously reported during Pi starvation (Fragoso *et al*., 2009; Trejo-Fregoso *et al*., 2022), we first sought to determine how Pi limitation affects canonical SnRK1 signalling *in vivo*. Previous evidence was largely derived from biochemical assays performed using protein extracts and artificial kinase substrates, which do not directly report SnRK1 signalling in its native cellular and metabolic context. We therefore used complementary approaches to assess SnRK1 activity during phosphate starvation. We first examined the phosphorylation status of Thr175 within the activation-loop (T-loop) of the SnRK1α catalytic subunits, a conserved regulatory residue whose phosphorylation is associated with kinase activation (Margalha *et al*., 2016). This analysis was performed in wild-type Col-0 seedlings following transfer from Pi-sufficient to Pi-deficient medium for 96 h. Pi starvation resulted in a marked reduction in the T-loop phosphorylation, whereas the abundance of the two major catalytic subunits, SnRK1α1/KIN10 and SnRK1α2/KIN11, remained largely unchanged (Fig. **1a**). Therefore, under these conditions, attenuation of T-loop phosphorylation occurred without an evident reduction in the abundance of either catalytic subunit. A similar reduction in T-loop phosphorylation was also observed in seedlings grown under Pi-deficient conditions from germination, indicating that attenuation of SnRK1 T-loop phosphorylation is not restricted to seedlings transferred to Pi-deficient medium (Fig. **S1a**). These results indicate that Pi starvation is consistently associated with reduced T-loop phosphorylation of SnRK1.

**Figure 1.**
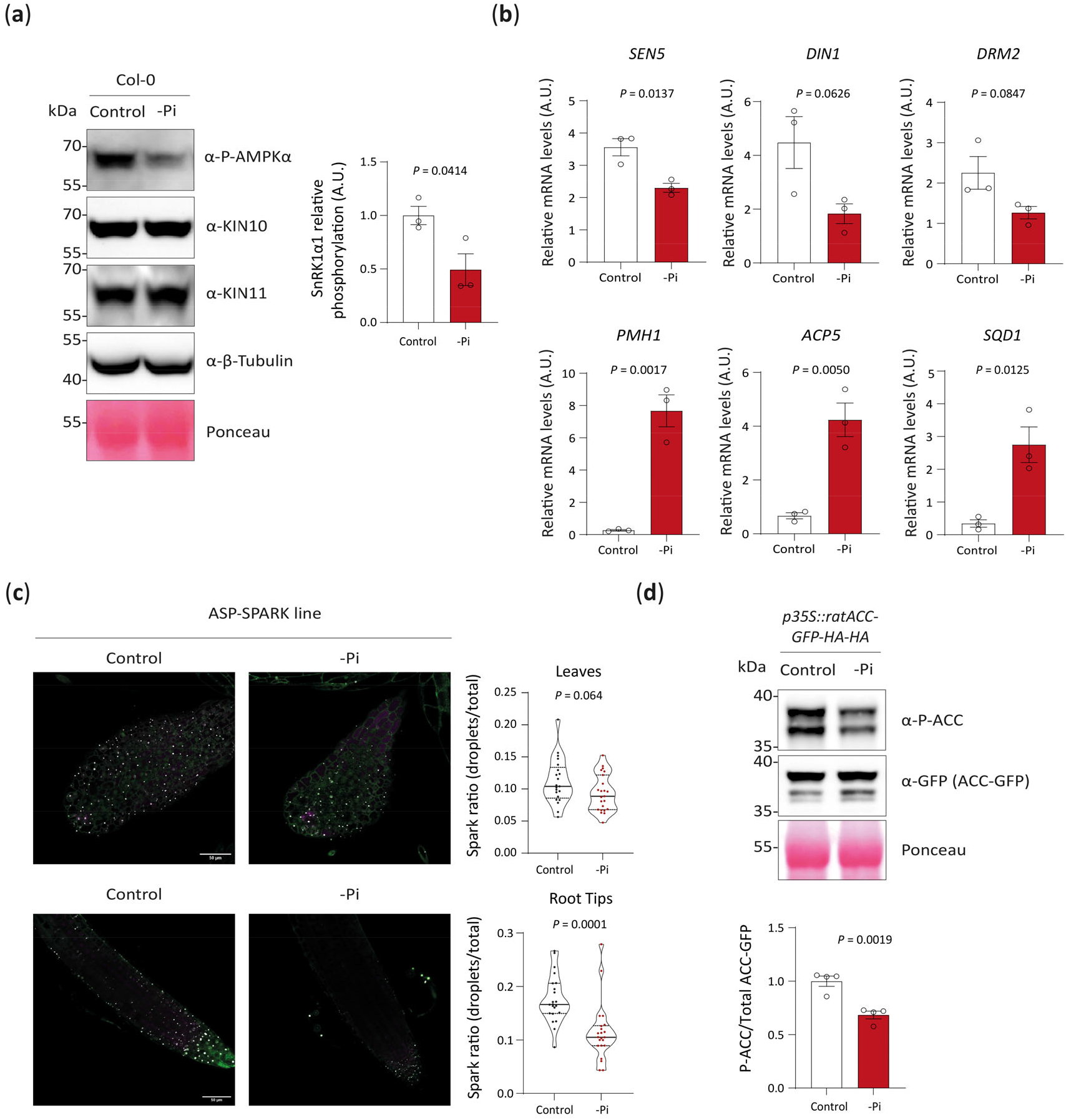
Phosphate starvation is associated with attenuation of SnRK1 activity *in vivo*. (a) Left, representative immunoblot analysis of SnRK1 activation-loop (T-loop) phosphorylation in Col-0 seedlings grown under Pi-sufficient (control) or Pi-deficient (-Pi) conditions for 96 h following transfer to the respective media. Membranes were probed with an antibody against phospho-AMPKα Thr172, which recognises the conserved T-loop phosphorylation sites of SnRK1α1/KIN10 (Thr175) and SnRK1α2/KIN11 (Thr176), and with antibodies against KIN10, KIN11 and β-Tubulin. β-Tubulin and Ponceau staining were used as loading controls. Right, quantification of SnRK1 T-loop phosphorylation. As the KIN10- and KIN11-associated phospho-AMPKα signals could not be reliably resolved, the combined phospho-AMPKα signal was normalised to the combined abundance of KIN10 and KIN11 [phospho-AMPKα signal (KIN10 + KIN11)/(KIN10 + KIN11 total)]. Data are presented as mean ± SEM, n = 3 biological replicates. *P*-values are indicated above the graphs, two-tailed Student’s *t-*test. (b) Expression of SnRK1-responsive genes (*SEN5*, *DIN1* and *DRM2*) and canonical phosphate-starvation-responsive genes (*PMH1*, *ACP5* and *SQD1*) in Col-0 seedlings grown under control or -Pi conditions for 96 h. Transcript levels were determined by RT-qPCR and normalised to *PDF2*. Data are presented as mean ± SEM, n = 3 biological replicates. *P*-values are indicated above the graphs, two-tailed Student’s *t-*test. (c) SnRK1 activity monitored using the genetically encoded ASP-SPARK reporter in root tips and leaves of seedlings grown under control or -Pi conditions for 6 days. Each dot represents an individual seedling. Data are presented as mean ± SEM, n = 21-23 seedlings. *P*-values are indicated, two-tailed Student’s *t-*test. Left, representative pictures obtained by confocal imaging. (d) Up, representative immunoblot analysis of phosphorylation of Ser79 in the ACC-GFP reporter in seedlings grown under control or -Pi conditions for 96 h. ACC Ser79 phosphorylation was detected using an anti-phospho-(S79)-ACC antibody, and ACC-GFP total abundance was assessed using an anti-GFP antibody. Ponceau staining was used as a loading control. Down, quantification of ACC phosphorylation [P(S79)-ACC/total ACC]. n = 3, *P*-values are indicated above the graphs, two-tailed Student’s *t-*test.

We next examined whether the reduction in SnRK1 T-loop phosphorylation was accompanied by changes in the expression of genes previously associated with SnRK1 activity (Baena-González *et al*., 2007). Following 96 h of Pi starvation, the expression of the SnRK1-responsive genes *DIN1*, *SEN5* and *DRM2* was reduced, whereas the canonical Pi-starvation-responsive genes *PMH1*, *ACP5* and *SQD1* were strongly induced (Bustos *et al*., 2010), confirming the establishment of the PSR under these conditions (Fig. **1b**). A similar reduction in SnRK1-responsive gene expression was observed after only 24 h of Pi starvation in liquid medium (Fig. **S1b**), indicating that this transcriptional response is reproducible across different experimental conditions and time scales.

We next sought to directly monitor SnRK1 activity *in vivo* using the genetically encoded ASP-SPARK reporter (Safi *et al*., 2023). In seedlings grown for 6 days under Pi-deficient conditions, Pi starvation resulted in a significant reduction in the SPARK ratio in root tips, consistent with decreased SnRK1 activity in these tissues (Fig. **1c**). A similar trend was observed in leaves, although the reduction did not reach statistical significance. Thus, direct monitoring of SnRK1 activity *in planta* supports a reduction in SnRK1 signalling during Pi starvation.

As an independent approach to assess SnRK1 activity *in vivo*, we used transgenic seedlings expressing the rat ACC-GFP reporter, in which phosphorylation of Ser79 provides a well-established readout of SnRK1 activity (Muralidhara *et al*., 2021). Pi starvation resulted in a marked reduction in ACC Ser79 phosphorylation, whereas the abundance of the ACC-GFP reporter remained largely unchanged (Fig. **1d**). A comparable reduction in ACC phosphorylation was also observed in seedlings grown under Pi-deficient conditions from germination for 6 days (Fig. **S1c**). Together, these complementary approaches consistently indicate that Pi starvation attenuates canonical SnRK1 signalling *in vivo* across distinct experimental regimes.

### Reducing SnRK1 activity enhances PSR-associated physiological responses, whereas increasing SnRK1 activity restrains them

Having established that Pi starvation is associated with attenuation of canonical SnRK1 signalling *in vivo*, we next asked whether this reduction in SnRK1 activity functionally contributes to plant adaptation to Pi limitation. To address this question, we examined physiological responses associated with the PSR in complementary genetic backgrounds with reduced or elevated SnRK1 activity. We first analysed primary root growth and anthocyanin accumulation in the single *snrk1α1* and *snrk1α2* mutants. However, neither genotype displayed clear differences from wild-type plants in the physiological responses examined, suggesting that partial functional redundancy between the two catalytic subunits may mask the effects of individual mutations (Fig. **S2a,b**) (Baena-González *et al*., 2007; Zacharaki *et al*., 2022).

We next used genetic backgrounds with stronger alterations in SnRK1 activity, including *sesquiα2-1*, a partial loss-of-function background with reduced SnRK1 activity (Belda-Palazón *et al*., 2020), and SnRK1α1-overexpressing (SnRK1α1-OE) plants with elevated SnRK1 activity (Jossier *et al*., 2009). The PSR regulator mutant *phr1-1* was included as a reference for impaired responses to Pi starvation (Rubio *et al*., 2001). Under Pi-deficient conditions, *phr1-1* displayed strongly reduced primary root growth and anthocyanin accumulation, whereas *sesquiα2-1* showed the opposite phenotype, with enhanced root growth and anthocyanin accumulation compared with the wild-type (Fig. **2a,b**). Conversely, SnRK1α1 overexpression exacerbated the inhibition of primary root growth while attenuating anthocyanin accumulation under Pi-deficient conditions (Fig. **2a,b**). Because the contrasting physiological phenotypes could potentially reflect differences in the Pi status of the plants rather than differences in their response to Pi starvation, we quantified soluble Pi levels under Pi-deficient conditions. Despite the marked differences in primary root growth and anthocyanin accumulation, soluble Pi levels were comparable among the genotypes, with the exception of *phr1-1*, which showed significantly lower soluble Pi levels (Fig. **2c**), indicating that these contrasting responses cannot be simply explained by differences in the bulk soluble Pi pool.

**Figure 2.**
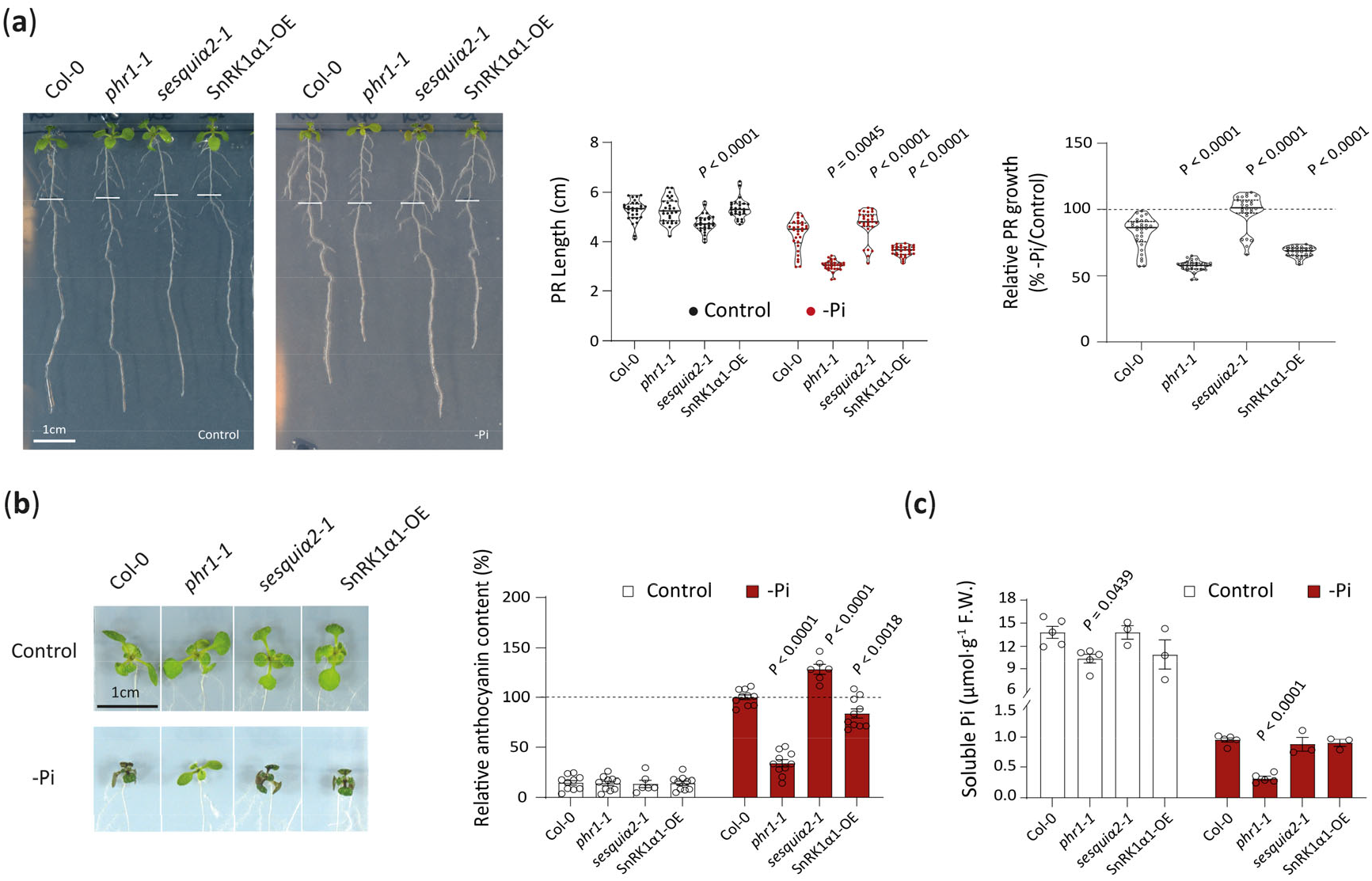
Altered SnRK1 activity modulates physiological responses to phosphate starvation. (a) Primary root (PR) growth of Col-0, *phr1-1*, *sesquiα2-1* and SnRK1α1-OE seedlings under Pi-sufficient (control) and Pi-deficient (-Pi) conditions. Left, representative images of seedlings after 6 days of vertical growth following transfer to control or -Pi medium. Scale bar, 1 cm. Middle, quantification of the PR length under control and -Pi conditions. Violin plots show the distribution of individual measurements, with each point representing an individual seedling. n = 28-30 seedlings per genotype and condition. *P*-values denote statistically significant differences from wild-type (Col-0) within the same treatment (two-way ANOVA followed by Dunnett’s multiple comparisons test). Right, quantification of relative PR growth under -Pi conditions, expressed relative to the corresponding control condition for each genotype (set to 100%). Violin plots show the distribution of individual measurements, with each point representing an individual seedling. *P*-values denote statistically significant differences from wild-type (Col-0) within the same treatment (One-way ANOVA followed by Dunnett’s multiple comparisons test). (b) Left, representative images of the seedlings grown under control or -Pi conditions for 7 days after transfer. Right, quantification of the relative anthocyanin content. Values are expressed relative to Col-0 under -Pi conditions (set to 100%). Data represent the mean ± SEM. n = 6-10 biological replicates. *P*-values denote statistically significant differences from wild-type (Col-0) within the same treatment (two-way ANOVA followed by Dunnett’s multiple comparisons test). (c) Soluble Pi content in seedlings grown under control or -Pi conditions for 6 days after transfer. Soluble Pi content is expressed as µmol Pi g⁻¹ FW. Data represent the mean ± SEM. n = 3-5 biological replicates. *P*-values denote statistically significant differences from wild-type (Col-0) within the same treatment (one-way ANOVA followed by Dunnett’s multiple comparisons test).

To further validate the relationship between SnRK1 activity and physiological responses to Pi starvation, we generated independent artificial microRNA (amiR) lines targeting both *SnRK1α1/KIN10* and *SnRK1α2/KIN11* (*amiR-KIN10/11-1, -2* and *-3*). To validate SnRK1 silencing, we analysed *KIN10* and *KIN11* transcript abundance by RT-PCR and observed reduced transcript levels in *amiR-KIN10/11-2* and *-3* compared with Col-0 (Fig. **S3a**). We next assessed the physiological consequences of reduced SnRK1 expression under carbon starvation. Following 7 days of growth under standard light conditions, seedlings were transferred to darkness for 7 days and subsequently allowed to recover under standard light conditions, in the presence or absence of 1% sucrose. The reduced *KIN10* and *KIN11* transcript abundance in *amiR-KIN10/11-2* and *-3* was associated with increased sensitivity to carbon starvation, as reflected by reduced seedling fresh weight, particularly in the absence of exogenous sucrose (Fig. **S3b**). Based on these results, *amiR-KIN10/11-2* and *-3* were selected for further analysis under Pi-deficient conditions. Under Pi-deficient conditions, both amiR lines showed a less pronounced inhibition of primary root growth than the corresponding control plants, together with enhanced anthocyanin accumulation (Fig. **S3c,d**). These results independently support an association between reduced SnRK1 expression and enhanced physiological responses to Pi starvation.

To determine whether the effects observed upon SnRK1α1 overexpression were specific to this catalytic subunit, we also generated independent SnRK1α2-OE lines, which showed increased KIN11 protein abundance, accompanied by a proportional increase in T-loop phosphorylation (Fig. **S4a**). Similar to SnRK1α1-overexpressing plants, both independent SnRK1α2-OE lines showed a more pronounced inhibition of primary root growth than the corresponding control plants under Pi-deficient conditions, together with reduced anthocyanin accumulation (Fig. **S4b,c**). Together, these complementary genetic approaches reveal an inverse relationship between SnRK1 activity and the magnitude of physiological responses associated with the PSR, with reduced SnRK1 activity enhancing, and elevated SnRK1 activity restraining, these responses under Pi limitation.

### SnRK1 controls the timing and magnitude of the proteomic response to phosphate starvation

To determine whether these contrasting physiological responses were reflected at the proteome level, we performed quantitative proteomic profiling of wild-type Col-0, SnRK1α1-OE, *amiR-KIN10/11-3* and *sesquiα2-1* seedlings exposed to Pi-sufficient or Pi-deficient conditions. Seedlings were harvested after 2 and 4 days of treatment, and shoots and roots were collected and analysed separately. The resulting datasets showed robust proteome coverage and good reproducibility across experimental groups (Table **S4**). Protein intensity distributions were broadly comparable among samples, most proteins were consistently quantified across biological replicates, and pairwise correlations indicated high reproducibility in both root and shoot datasets (Fig. **S5** and **S6**). Principal component analysis (PCA) revealed a progressive restructuring of the proteome in response to Pi availability (Fig. **S7a**). At 2 days, proteomic profiles from Pi-sufficient and Pi-deficient seedlings showed substantial overlap in both roots and shoots. By contrast, after 4 days of treatment, the PCA revealed an emerging separation between Pi-sufficient and Pi-deficient samples, with this pattern being more pronounced in shoots. Notably, samples from the different genotypes showed substantial overlap within the global PCA structure, indicating that altered SnRK1 activity did not represent the dominant source of proteome-wide variation. Nevertheless, genotype-dependent differences became more apparent after prolonged Pi deprivation, particularly in shoots. These observations suggested that altered SnRK1 activity may primarily affect the kinetics and magnitude of specific proteomic responses to Pi deprivation, rather than causing an extensive reorganisation of the proteome.

To characterize the proteomic response to Pi deprivation, we first performed differential abundance analyses comparing Pi-deficient with Pi-sufficient conditions within each genotype at 2 and 4 days of treatment (Table **S5**). The extent of proteome remodelling varied markedly among genotypes, tissues and time points (Fig. **S7b**). Notably, some genotypes, mainly the Col-0 and the *sesquiα2-1*, showed pronounced changes in protein abundance, particularly after 4 days of Pi deprivation, whereas more limited responses were observed in the other genotype-tissue combinations. Functional enrichment analysis of proteins differentially abundant between phosphate deficiency and control conditions in each genotype revealed associations with nutrient starvation, phosphate homeostasis and related metabolic and molecular processes (Table **S6** and Fig. **S7c**). To examine how altered SnRK1 activity influenced the proteomic response under Pi starvation, we next performed differential abundance analyses comparing each SnRK1-altered genotype with wild-type plants under Pi-deficient conditions at 2 and 4 days of treatment (Table **S5**). Volcano plots illustrated the magnitude and direction of these genotype-dependent differences, revealing distinct patterns of protein accumulation and depletion across genotypes, tissues and time points (Fig. **S8**).

To delineate how SnRK1 interplays with the response to Pi starvation, we focused on a curated set of PSR-associated proteins (Bustos *et al*., 2010) and examined how their abundance was affected in genotypes with altered SnRK1 activity across tissues and treatment duration (Fig. **3**). The abundance patterns of the SnRK1 catalytic subunits KIN10 and KIN11 were consistent with the expected SnRK1 genetic backgrounds of the different lines, while SnRK1 activity-associated proteins showed genotype-dependent patterns consistent with altered SnRK1 status. Importantly, Pi deprivation induced coordinated changes in proteins involved in Pi signalling, phosphate recycling, lipid remodelling, flavonoid/anthocyanin metabolism and general stress responses, with the magnitude of these changes varying across time points, SnRK1 genotypes and tissues (Figs. **3**, **S9** and **S10**). In shoots, genotype-dependent differences were already evident under Pi-sufficient conditions for several metabolic and stress-related proteins, particularly in the *amiR-KIN10/11-3* and *sesquiα2-1* backgrounds. Upon Pi deprivation, PSR-associated proteins progressively accumulated in Col-0 during Pi deprivation, with this response occurring earlier in the *amiR-KIN10/11-3* and *sesquiα2-1* backgrounds, whereas their accumulation was delayed in SnRK1α1-OE. Moreover, under Pi starvation, the SnRK1 partial loss-of-function backgrounds showed higher abundance of proteins involved in Pi sensing and transport, phosphate recycling, lipid remodelling, flavonoid/anthocyanin metabolism and general stress responses than Col-0 (Figs. **3a** and **S9**). In roots, genotype-dependent differences in PSR-associated proteins were most evident after 2 days of Pi deprivation. SnRK1α1-OE showed reduced abundance of several PSR-associated proteins relative to Col-0, whereas *sesquiα2-1* showed higher abundance of several of these proteins at this early time point. By 4 days of Pi deprivation, these genotype-dependent differences were less pronounced, with PSR-associated protein abundance becoming more similar across genotypes (Figs. **3b** and **S10**). Thus, altered SnRK1 activity was associated with differences in both the magnitude and timing of PSR-associated proteome remodelling, with reduced SnRK1 activity generally associated with an earlier and/or stronger accumulation of PSR-associated proteins.

**Figure 3.**
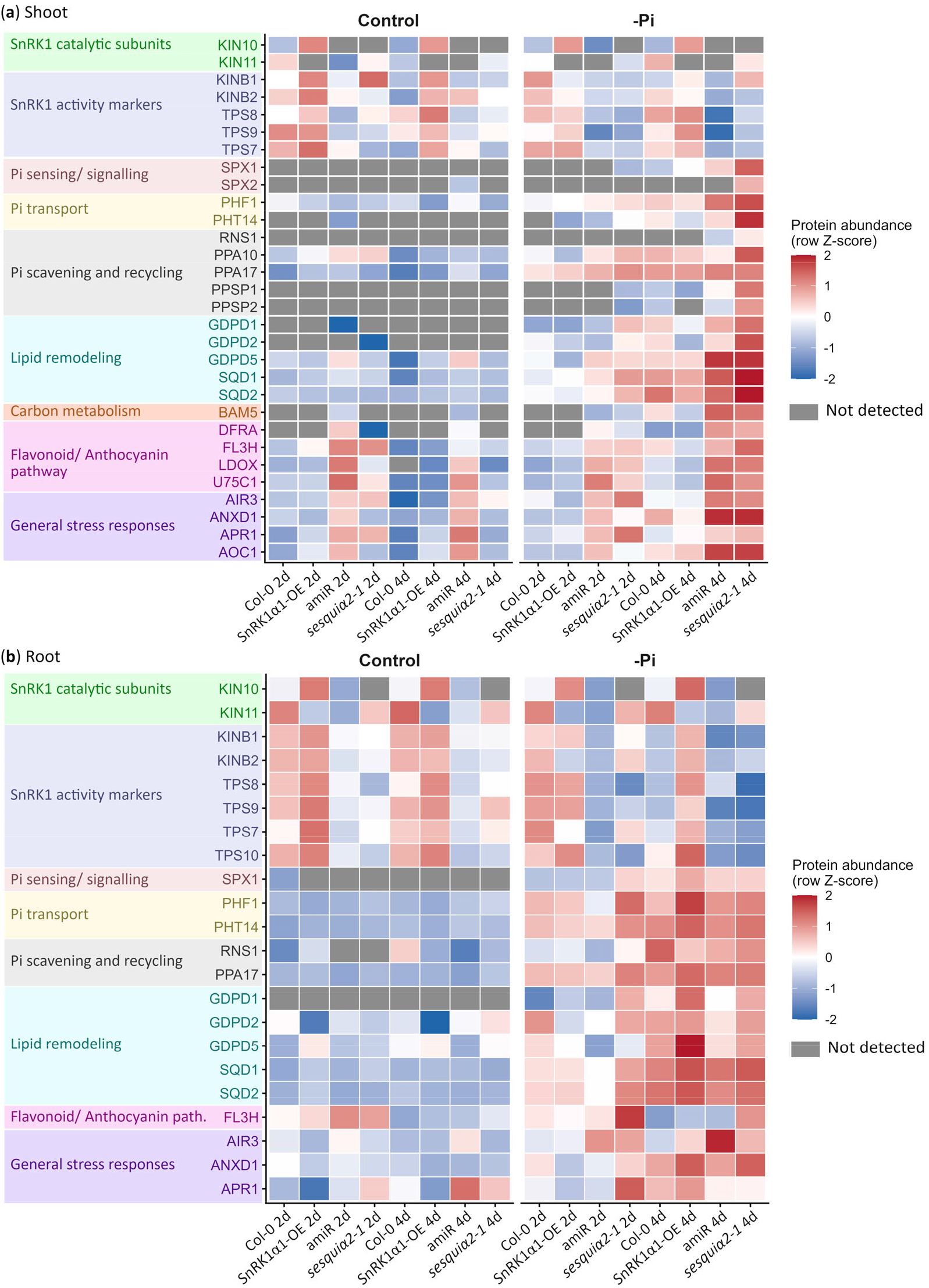
SnRK1 controls the timing and magnitude of the proteomic response to phosphate starvation. (a) Heatmap showing the abundance of selected PSR-associated proteins in shoots of Col-0, SnRK1α1-OE, *amiR-KIN10/11-3* (amiR) and *sesquiα2-1* seedlings grown under Pi-sufficient (control) or Pi-deficient (-Pi) conditions for 2 or 4 days. (b) Corresponding heatmap for roots. Proteins are grouped according to their functional association with SnRK1 signalling, Pi sensing/signalling, Pi transport, Pi scavenging and recycling, lipid remodelling, carbon metabolism, flavonoid/anthocyanin metabolism and general stress responses. Values represent the mean row-wise Z-score of three biological replicates for each genotype, condition and time point. Red and blue indicate relatively higher and lower protein abundance, respectively. Grey indicates proteins that were not detected.

### SnRK1 shapes metabolic adaptation to phosphate starvation

To determine whether these proteomic changes were accompanied by alterations in primary metabolism, we next analysed the abundance of primary metabolites in Col-0, SnRK1α1-OE and *sesquiα2-1* seedlings cultivated as for the proteome assay (Table **S7**). PCAs showed a clear separation of metabolite profiles according to Pi availability in both shoots and roots, with Pi status accounting for a major component of the variation captured by PC1 (58.1% and 65.6% in shoots and roots, respectively). Within Pi-deficient samples, treatment duration contributed to a further separation along PC2, consistent with continued metabolic remodelling during Pi deprivation. Genotype-dependent separation was also observed within Pi-deficient samples, although the extent of separation differed between shoots and roots (Fig. **S11a**). PCA variable plots indicated that this separation was largely associated with opposing patterns of hexose-phosphate and glycerol-3-phosphate pools versus soluble sugars, amino acids and organic acids in both tissues (Fig. **S11b**). Thus, Pi starvation induced earlier and more SnRK1-activity-dependent changes in the metabolome than in the proteome.

We therefore next examined how SnRK1 perturbation affected the abundance of individual metabolites across genotypes, tissues and treatment duration under Pi-sufficient and Pi-deficient conditions (Table **S8** and Fig. **4**). Across genotypes, Pi deprivation induced a broad metabolic response consistent with established features of phosphate starvation (Dissanayaka et al., 2021; Roychowdhury et al., 2023). Phosphorylated carbohydrate intermediates, including glucose-6-phosphate, fructose-6-phosphate and glycerol-3-phosphate, generally decreased under -Pi conditions, whereas soluble sugars and organic acids accumulated. The response was particularly pronounced for organic acids and amino acids in roots, while metabolic changes in shoots became more extensive at the later time point. Thus, the experimental conditions triggered a robust metabolic response to Pi deprivation across both tissues.

**Figure 4.**
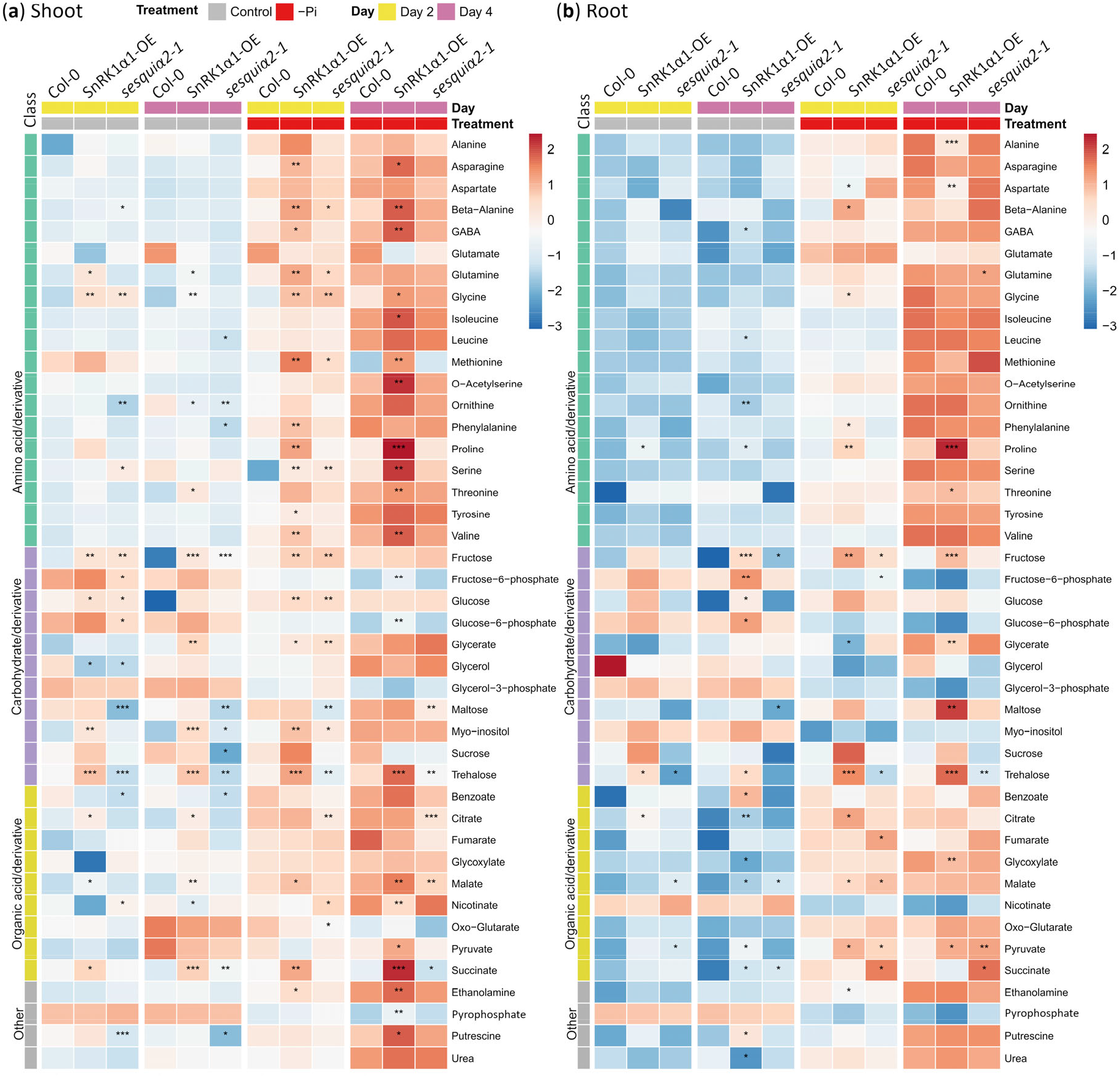
SnRK1 shapes metabolic adaptation to phosphate starvation. (a) Heatmap showing primary metabolite relative abundance in shoots of Col-0, SnRK1α1-OE and *sesquiα2-1* seedlings grown under Pi-sufficient (control) or Pi-deficient (-Pi) conditions for 2 or 4 d. (b) Corresponding heatmap for roots. Metabolites are grouped according to their major metabolic classes. Values represent the mean relative abundance of five biological replicates and are shown as row-wise Z-scores calculated across the samples included in each heatmap. Red and blue indicate relatively higher and lower metabolite abundance, respectively. Asterisks indicate significant differences between each SnRK1-altered genotype and Col-0 under the corresponding treatment and time point, determined by Welch’s *t*-test followed by Benjamini-Hochberg correction for multiple testing. *, *P* ≤ 0.05; **, *P* ≤ 0.01; and ***, *P* **≤** 0.001.

To further examine genotype-dependent changes in primary metabolism under Pi deprivation, we next analysed metabolites that differed between each SnRK1-altered genotype and Col-0 under -Pi conditions. In shoots, altered SnRK1 activity was associated with metabolite-specific differences in basal metabolism under Pi-sufficient conditions, particularly within carbohydrate pools. Upon Pi deprivation, these differences were further reshaped, revealing distinct genotype-dependent responses. Wild-type Col-0 showed a coordinated metabolic adaptation to Pi deprivation, characterized by changes in amino-acid, carbohydrate and organic-acid metabolism. Increasing SnRK1 activity markedly altered this response, with the SnRK1α1-OE line showing stronger accumulation of amino acids and carbohydrates, together with attenuated depletion of hexose-phosphate metabolites and more pronounced changes in organic-acid abundance. In contrast, *sesquiα2-1* showed an attenuated metabolic response, with weaker amino-acid accumulation and reduced levels of selected carbohydrate and organic-acid metabolites, compatible with a reduced extent of metabolic reprogramming. Thus, in shoots, SnRK1 activity primarily influenced the magnitude and composition of metabolic reprogramming during Pi deprivation (Fig. **4a**). The enhanced metabolic response in SnRK1α1-OE and its attenuation in *sesquiα2-1* are in agreement with the established role of SnRK1 in coordinating metabolic reprogramming in response to energy limitation (Peixoto & Baena-González, 2022).

However, the situation was different in roots. Here, SnRK1 activity did not simply influence the magnitude of the response, but was associated with distinct metabolic configurations. Under Pi-sufficient conditions, the three genotypes showed relatively modest and metabolite-specific differences in basal metabolite abundance. Upon Pi deprivation, these differences became more pronounced and revealed distinct genotype-dependent responses. The wild-type Col-0 showed a coordinated metabolic response to Pi deprivation, including carbohydrate adjustments, depletion of hexose-phosphate metabolites and remodelling of organic acids. Increasing SnRK1 activity was associated with strong accumulation of soluble carbohydrates, particularly maltose, sucrose and trehalose, together with moderate organic-acid changes. This carbohydrate accumulation was not accompanied by a comparable increase in the organic-acid pools examined and was not accompanied by a corresponding enhancement of root growth in SnRK1α1-OE (Fig. **2a**). In contrast, *sesquiα2-1* did not show the strong carbohydrate accumulation observed in SnRK1α1-OE, but instead showed a more pronounced accumulation of selected organic acids, including pyruvate, succinate, fumarate and malate, particularly at the early stage of Pi deprivation (Fig. **4b**). This distinct metabolic profile was associated with the enhanced root growth observed in *sesquiα2-1* under Pi deprivation (Fig. **2a**). Together, these results indicate that SnRK1 contributes to metabolic homeostasis and shapes both the magnitude and the configuration of the metabolic response to phosphate starvation in a tissue-dependent manner.

## Discussion

Phosphate starvation requires extensive reprogramming of plant growth, metabolism and resource allocation, yet how this response is coordinated with cellular energy status remains incompletely understood. Here, we identify SnRK1 as an important regulator of this coordination. We show that phosphate deprivation is accompanied by a reduction in SnRK1 activity and T-loop phosphorylation, and that perturbing SnRK1 activity alters the timing and magnitude of the phosphate starvation response at physiological, proteomic and metabolic levels. Reduced SnRK1 activity was generally associated with an earlier and stronger deployment of phosphate-starvation-associated responses, whereas increased SnRK1 activity delayed or attenuated several components of the response. These effects were dependent on tissue and treatment duration and were accompanied by distinct metabolic configurations. Thus, our findings indicate that SnRK1 activity quantitatively tunes the extent and timing of the phosphate starvation response. We propose that this quantitative regulation allows plants to balance the metabolic investment required for phosphate-starvation adaptation with the maintenance of growth.

An important question arising from these findings is why SnRK1 activity is attenuated during phosphate starvation. Previous studies have revealed apparently divergent regulation of SnRK1 under Pi deficiency, suggesting that its regulation is more complex than a simple on-off response, with changes in catalytic subunit composition and activity that are not necessarily accompanied by changes in T-loop phosphorylation (Fragoso *et al*., 2009; Trejo-Fregoso *et al*., 2022). In contrast, under our experimental conditions, we detected a reduction in the activation T-loop phosphorylation status of SnRK1 catalytic subunits during phosphate starvation, without evidence for a corresponding change in their total abundance. Moreover, SnRK1α1- and SnRK1α2-overexpressing plants displayed similar physiological phenotypes, consistent with partially overlapping contributions of the two catalytic subunits to the regulation of the PSR. Conversely, Trejo-Fregoso *et al*. (2022) showed that SnRK1 can directly phosphorylate components of the PSR, including the central transcriptional regulator PHR1, and proposed that phosphorylation of PHR1 at Ser11 negatively regulates its transcriptional activity. These findings raise the possibility that attenuation of SnRK1 activity itself may facilitate full deployment of the PSR. This interpretation is consistent with our physiological analyses, which showed that reduced SnRK1 activity was associated with enhanced phosphate-starvation responses, whereas increased SnRK1 activity tended to attenuate these responses. Although T-loop phosphorylation is required for SnRK1 catalytic activity, its relationship with kinase activity *in vivo* appears to vary depending on the physiological context (Margalha *et al*., 2019). T-loop phosphorylation can be dynamically regulated under specific stress conditions, whereas in other contexts changes in T-loop phosphorylation do not necessarily parallel changes in SnRK1 activity. This apparent uncoupling is also relevant to the proposed default-active state of plant SnRK1, in which the catalytic subunits can retain activity independently of canonical regulatory subunits (Ramon *et al*., 2019). Against this background, the concomitant reduction in SnRK1 activity and T-loop phosphorylation observed here suggests that regulation of the T-loop phosphorylation state may contribute to SnRK1 attenuation during Pi starvation.

Phosphate starvation is not a passive response to nutrient limitation, but an active and metabolically demanding program. Adaptation to low Pi requires coordinated changes in phosphate acquisition and recycling, metabolic reprogramming, membrane lipid remodelling, root architecture and secondary metabolism, all of which require substantial carbon, energy and reducing power. These demands imply that deployment of the PSR must be coordinated with the metabolic capacity of the plant. Our physiological, proteomic and metabolomic data support a model in which SnRK1 acts as a quantitative metabolic brake on the PSR. In this context, the attenuation of SnRK1 activity during Pi starvation may increase metabolic flexibility by relaxing constraints on the allocation of cellular resources, thereby allowing greater investment in the processes required for adaptation to Pi limitation. Conversely, when SnRK1 activity is maintained or increased, this metabolic constraint may remain stronger, limiting investment in energetically costly components of the PSR. Importantly, greater metabolic investment in the PSR does not necessarily translate into enhanced growth. This was particularly evident in roots, where increased SnRK1 activity favoured a carbohydrate-rich metabolic state without a corresponding increase in root growth, whereas reduced SnRK1 activity was associated with metabolic changes that coincided with enhanced root growth. Thus, the physiological outcome of metabolic reprogramming under Pi limitation depends not only on the extent of the response, but also on how metabolic resources are allocated between adaptation and growth.

Interestingly, the contrasting effects of SnRK1 activity on primary root growth were not accompanied by differences in bulk soluble Pi content. Thus, the enhanced root growth of *sesquiα2-1* and the stronger inhibition of root growth in SnRK1α1-OE under Pi deficiency cannot be readily attributed to differences in the overall soluble Pi status of the seedlings. This does not necessarily imply equivalent Pi homeostasis, as steady-state Pi levels reflect the balance between Pi acquisition, utilization, remobilization and storage, while bulk measurements do not resolve the distribution of Pi between subcellular pools (Yang *et al*., 2017; Versaw & Garcia, 2017). Differences in Pi fluxes or compartmentalization could therefore remain undetected in our measurements. Indeed, previous studies have shown that altered intracellular Pi compartmentation can enhance Pi-starvation responses despite unchanged total Pi content (Hassler *et al*., 2012; Cardona-López *et al*., 2015). Moreover, Pi-starvation responses are regulated by local and systemic sensing and signalling mechanisms that are not necessarily determined solely by bulk tissue Pi levels. Consistent with this view, previous work has shown that Pi content can be uncoupled from physiological and molecular responses to Pi deficiency (Rouached *et al*., 2011). Importantly, our metabolomic analyses revealed distinct metabolic configurations in the SnRK1-altered genotypes despite their similar bulk soluble Pi content. Under Pi deficiency, SnRK1α1-OE roots accumulated soluble carbohydrates, including maltose, sucrose and trehalose, whereas *sesquiα2-1* roots showed stronger accumulation of selected organic acids, including pyruvate, succinate, fumarate and malate, concomitant with enhanced root growth. These observations suggest that SnRK1 influences how metabolic resources are configured and deployed in response to a given Pi status, rather than simply determining bulk cellular Pi content.

This balance between adaptation and growth is particularly relevant to the SnRK1-TOR regulatory axis. SnRK1 and TOR are generally viewed as antagonistic regulators of cellular resource allocation, with SnRK1 promoting metabolic restraint under conditions of limited energy and TOR promoting growth when resources are sufficient (Margalha *et al*., 2019). However, emerging evidence suggests that this antagonism is not absolute under Pi starvation. Recent work showed that TOR activity increases under Pi limitation and contributes to PSR gene induction through PHR1 stabilization, indicating that TOR can support adaptive responses under nutrient limitation rather than simply promoting growth (Awasthi *et al*., 2024). In this context, the attenuation of SnRK1 activity observed here may complement TOR activation by increasing metabolic flexibility and allowing greater resource allocation towards adaptation, while the balance between SnRK1 and TOR activity may influence how these resources are ultimately distributed between adaptation and growth. Thus, rather than acting as strictly opposing switches, SnRK1 and TOR may function as interconnected regulators that coordinate metabolic status, adaptation and growth during Pi limitation.

Together, our findings suggest that Pi starvation-induced attenuation of SnRK1 releases a quantitative metabolic brake, increasing the flexibility with which plants balance resource allocation between phosphate-starvation adaptation and growth.

## Acknowledgements

We thank Filip Rolland, Javier Paz-Ares and Jelle Van Leene for the *p35S::ratACC-GFP-HA-HA*, *phr1-1,* and *ASP-SPARK A. thaliana* transgenic lines, respectively. The proteomic analysis was performed in the proteomics facility of SCSIE University of Valencia. This proteomics laboratory is a member of Proteored. The authors acknowledge the assistance of the IBMCP Scientific and Technical Services, including the Advanced Microscopy Unit, the Plant Growth Facility, for their support during the course of this work. We gratefully acknowledge the financial support that made this study possible. This research was funded the CIDEGENT program of excellence of GVA, grant number CIDEXG/2022/27 (awarded to B.B.-P.), and the Spanish Ministry of Science, Innovation and Universities (MCIU), grant numbers PID2022-142412NB-I00 (awarded to B.B.-P. and A.F.), PID2024-162615OB-I00 and RYC2022-037020-I (A.G.-M.) and CNS2024-154566 (awarded to B.B.-P.). A.V.-F. was the beneficiary of PhD contract CIDEXG/2022/27-02, funded by GVA. A.C.-A. was the beneficiary of PhD contract PRE2022-105204, funded by MCIN/AEI/10.13039/501100011033 and by the European Social Fund Plus (FSE+). We acknowledge financial support from the institutional grant CEX2025-001626-S and CEX2019-000902-S (Center of Excellence Severo Ochoa) funded by MCIU/AEI/10.13039/501100011033 and the CERCA Program/Generalitat de Catalunya. We acknowledge FCT - Fundação para a Ciência e a Tecnologia, I.P., through Green-it Bioresources for Sustainability R&D Unit (UID/04551/2025, DOI: 10.54499/UID/04551/2025; UID/PRR/04551/2025, DOI: 10.54499/UID/PRR/04551/2025), LS4FUTURE Associated Laboratory (LA/P/0087/2020, DOI: 10.54499/LA/P/0087/2020), grant 2024.17074.PEX (DOI: 10.54499/2024.17074.PEX), and individual contract (2023.11076.TENURE.104) to A.C. We acknowledge the use of OpenAI’s ChatGPT to assist with language editing and refinement of the manuscript. The authors take full responsibility for the scientific content, interpretation and conclusions presented in this work.

## Author contributions

A.C.-A., A.V.-F., A.A., A.C., L.F., C.C.-R., and B.B.-P. conducted experiments. A.A., A.R.F and A.G.-M provided expertise in metabolomic and proteomic analyses and R-based data analysis. E.B.-G. and A.F. provided tools and expertise on SnRK1 signalling and molecular integration, actively supporting the conceptual work. B.B.-P. conceived and designed the project. B.B.-P. directed and supervised all research activities and prepared figures. B.B.-P. wrote the manuscript. A.A., A.C., A.R.F., E.B.-G., and A.F. reviewed the manuscript. All authors have read and agreed to the published version of the manuscript.

## Data availability

All data supporting the findings of this study are available within the article and its Supporting Information. Additional data and materials related to this study are available from the corresponding author upon reasonable request. The mass spectrometry proteomics data have been deposited to the ProteomeXchange Consortium via the PRIDE (Perez-Riverol *et al*., 2025) partner repository with the dataset identifier PXD085154.

