## Supplemental info for "Phosphate Deficiency Attenuates SnRK1 Activity to Modulate the Phosphate Starvation Response in *Arabidopsis thaliana*"

The following Supporting Information is available for this article:

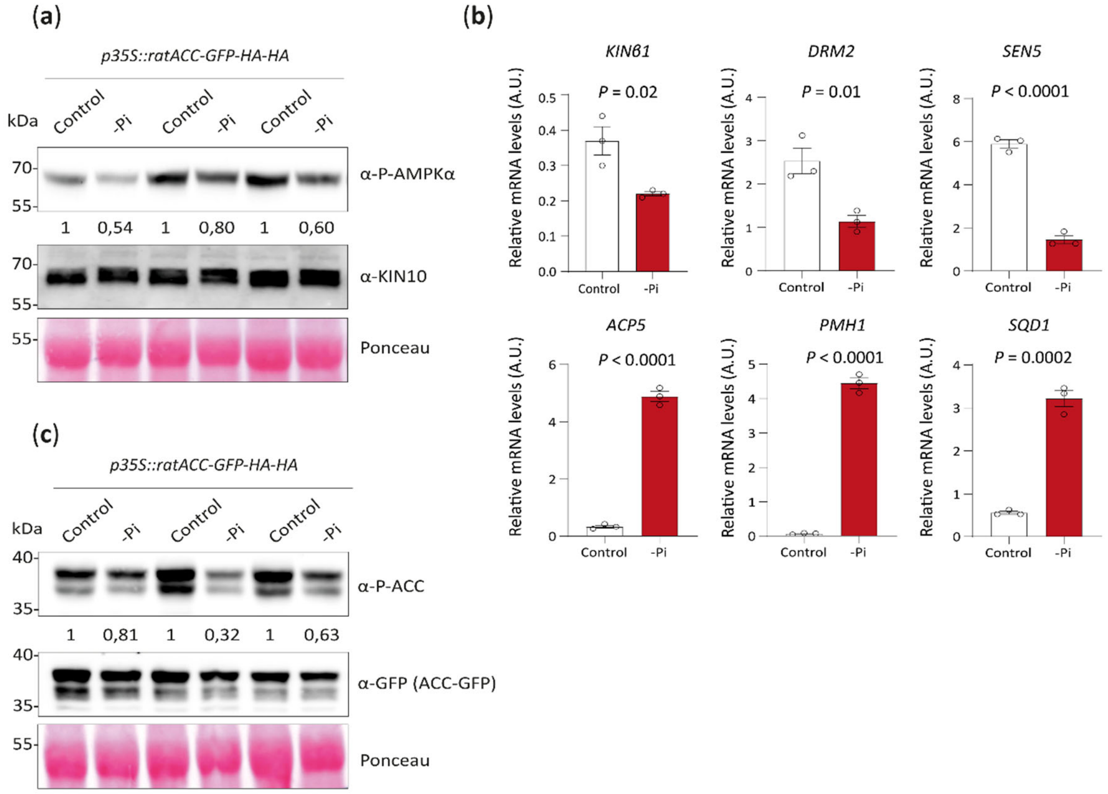


**Figure S1. Phosphate starvation attenuates SnRK1 signalling under independent experimental conditions.** (a) Immunoblot analysis of SnRK1 activation-loop (T-loop) phosphorylation in *p35S::ratACC-GFP-HA-HA* seedlings grown under Pi-sufficient (control) or Pi-deficient (-Pi) conditions for 6 days from germination. Membranes were probed with an antibody against phospho-AMPKα Thr172, which recognises the conserved T-loop phosphorylation sites of SnRK1α1/KIN10 (Thr175) and with an antibody against KIN10. Ponceau staining was used as a loading control.

Numbers indicate the relative phospho-AMPKα signal normalised to KIN10 abundance, with each -Pi sample normalised to its corresponding +Pi control (set to 1). (b) Expression of the SnRK1-responsive genes *KINB1*, *DRM2* and *SEN5*, and the phosphate-starvation-responsive genes *ACP5*, *PMH1* and *SQD1*, in seedlings subjected to Pi starvation for 24 h in liquid medium. Transcript levels were determined by RT-qPCR and normalised to *PDF2*. Data are presented as mean ± SEM (n = 3 biological replicates). *P*-values are indicated above the graphs, two-tailed Student´s *t*-test. (c) Representative immunoblot analysis of ACC Ser79 phosphorylation in *p35S::ratACC-GFP-HA-HA* seedlings grown under Pi-sufficient (control) or Pi-deficient (-Pi) conditions for 6 days from germination. Membranes were probed with an anti-phospho-(S79)-ACC antibody and an anti-GFP antibody to detect phosphorylated and total ACC-GFP, respectively. Ponceau staining was used as a loading control. Numbers below the blots indicate the relative phospho-ACC signal normalised to ACC-GFP abundance, with each -Pi sample normalised to its corresponding +Pi control (set to 1).

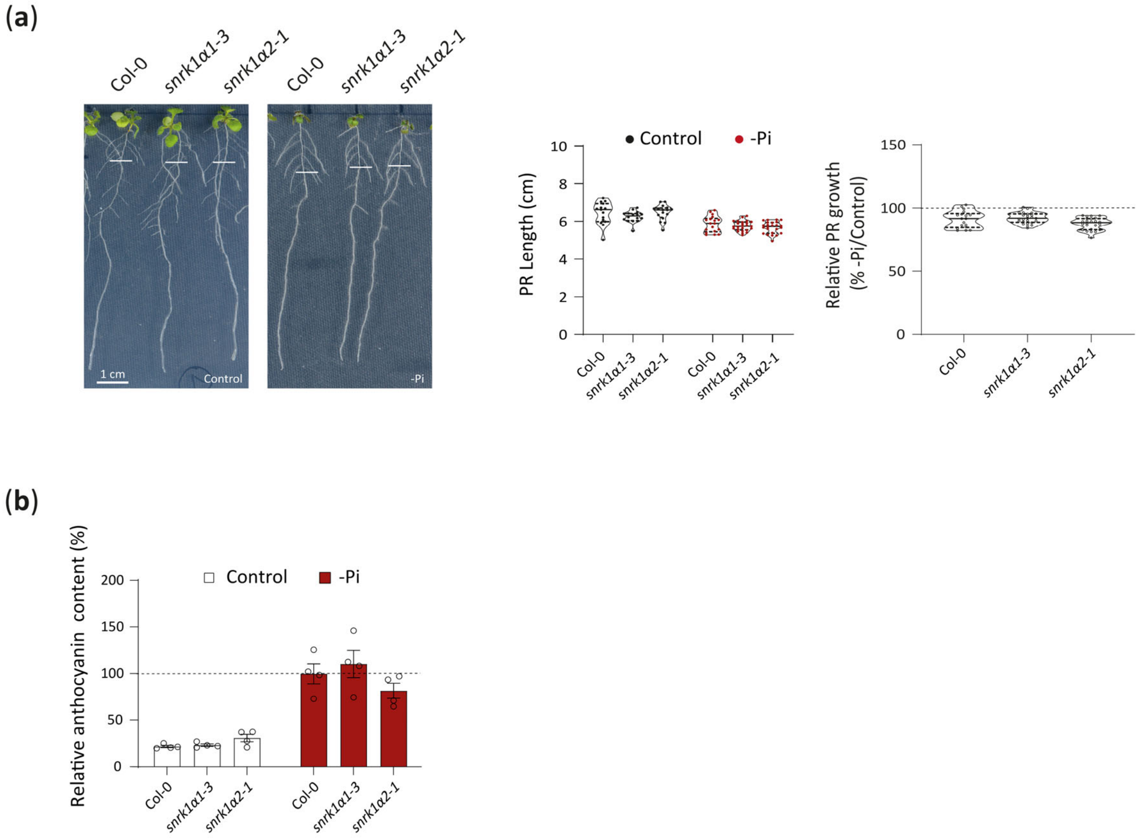


**Figure S2. Physiological responses of *snrk1α1-3* and *snrk1α2-1* mutants to phosphate starvation.** (a) Primary root growth (PR) of Col-0, *snrk1α1-3* and *snrk1α2-1* seedlings under Pi-sufficient (control) and Pi-deficient (-Pi) conditions. Left, representative images of seedlings after 6 days of vertical growth following transfer to control or -Pi medium. Scale bar, 1 cm. Middle, quantification of PR length. n = 15-25 seedlings per genotype and condition. Right, quantification of relative PR growth under -Pi conditions, expressed relative to the corresponding control condition for each genotype (set to 100%). Violin plots show the distribution of individual measurements, with each point representing an individual seedling. The absence of a *P*-value indicates a lack of statistically significant differences from wild-type (Col-0) within the same treatment. Two-way (left graph) or one-way (right graph) ANOVA followed by Dunnett's multiple comparisons test. (b) Relative anthocyanin content in Col-0, *snrk1α1-3* and *snrk1α2-1* seedlings grown under control or -Pi conditions for 6 days following transfer to control or -Pi medium. Values are expressed relative to Col-0 under -Pi conditions (set to 100%). Data represent the mean ± SEM. n = 4 biological replicates. The absence of a *P*-value indicates a lack of statistically significant differences from wild-type (Col-0) within the same treatment (two-way ANOVA followed by Dunnett's multiple comparisons test).

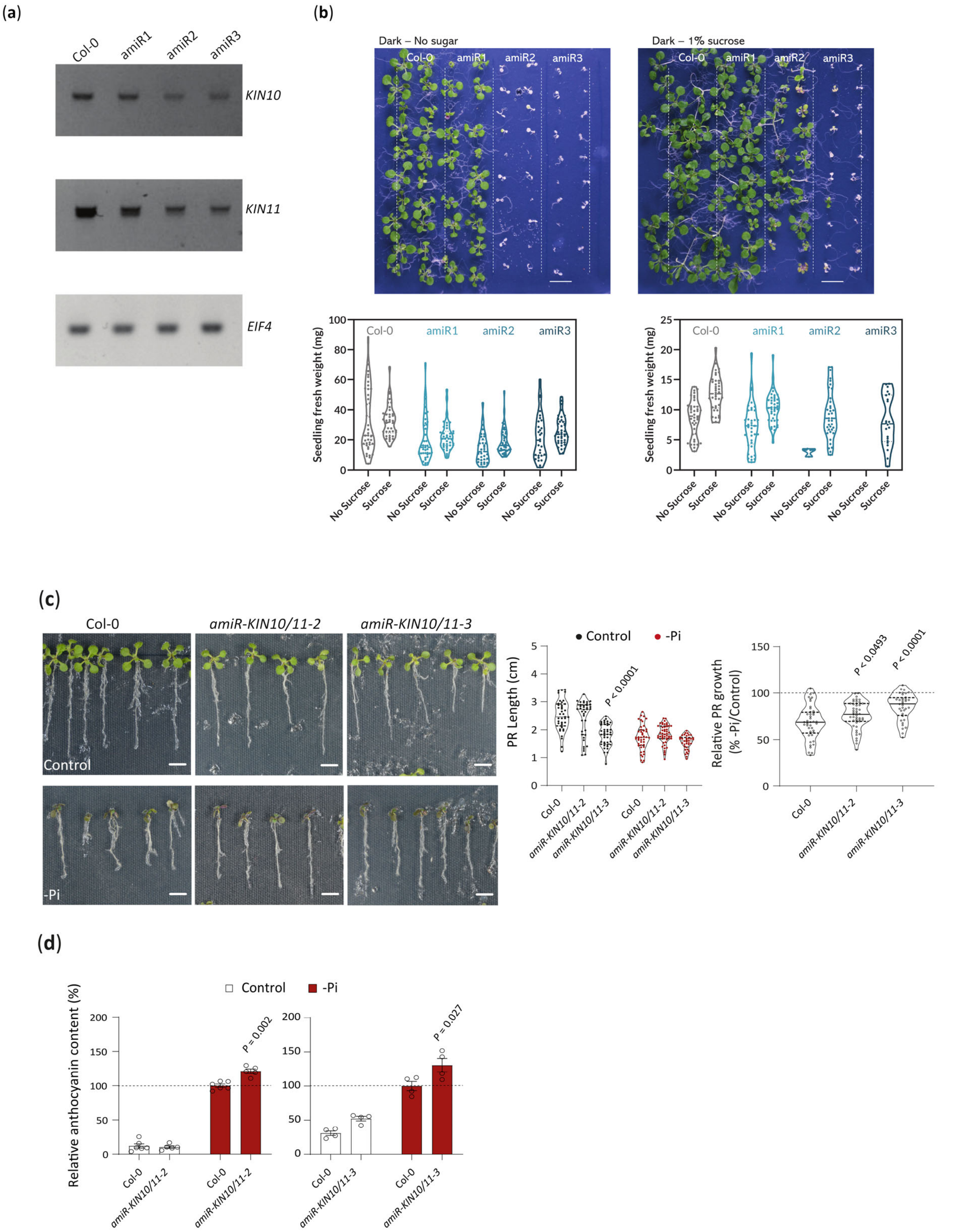


**Figure S3. Reduced SnRK1α1/2 expression in *amiR-KIN10/11* lines attenuates the inhibition of root growth and**

**enhances anthocyanin accumulation during phosphate starvation.** (a) Representative RT-PCR analysis of *KIN10* and *KIN11* transcript abundance in Col-0 and three independent *amiR-KIN10/11* lines (amiR1, amiR2 and amiR3). *EIF4* was used as a control for transcript abundance. (b) Seedling fresh weight of Col-0 and the three independent *amiR-KIN10/11* lines grown under 16 h light/8 h dark cycles for 7 days, followed by 7 days in darkness and a subsequent 7 days recovery period under 16 h light/8 h dark cycles. Seedlings were grown on medium without sucrose or supplemented with 1% sucrose. Upper panels, representative images of seedlings. Lower panels, quantification of seedling fresh weight. Violin plots show the distribution of individual measurements, with each point representing an individual seedling. (c) Primary root growth of Col-0, *amiR-KIN10/11-2* and *amiR-KIN10/11-3* seedlings under Pi-sufficient (control) and Pi-deficient (-Pi) conditions. Left, representative images of seedlings after 6 days of horizontal growth after transfer to control or -Pi medium. Scale bar, 1 cm. Middle, quantification of PR length. Right, quantification of PR growth under -Pi conditions, expressed relative to the corresponding control condition for each genotype (set to 100%). Violin plots show the distribution of individual measurements, with each point representing an individual seedling. n = 36-49 seedlings per genotype and condition. *P*-values denote statistically significant differences from wild-type (Col-0) within the same treatment. Two-way (left graph) or one-way (right graph) ANOVA followed by Dunnett's multiple comparisons test. (c) Relative anthocyanin content in Col-0, *amiR-KIN10/11-2* and *amiR-KIN10/11-3* seedlings grown under control or -Pi conditions for 96 h following transfer to control or -Pi medium. Values are expressed relative to Col-0 under -Pi conditions (set to 100%). Data represent the mean ± SEM. n = 4 biological replicates. *P*-values denote statistically significant differences from wild-type (Col-0) within the same treatment. Two-way ANOVA followed by Tukey's multiple comparisons test.

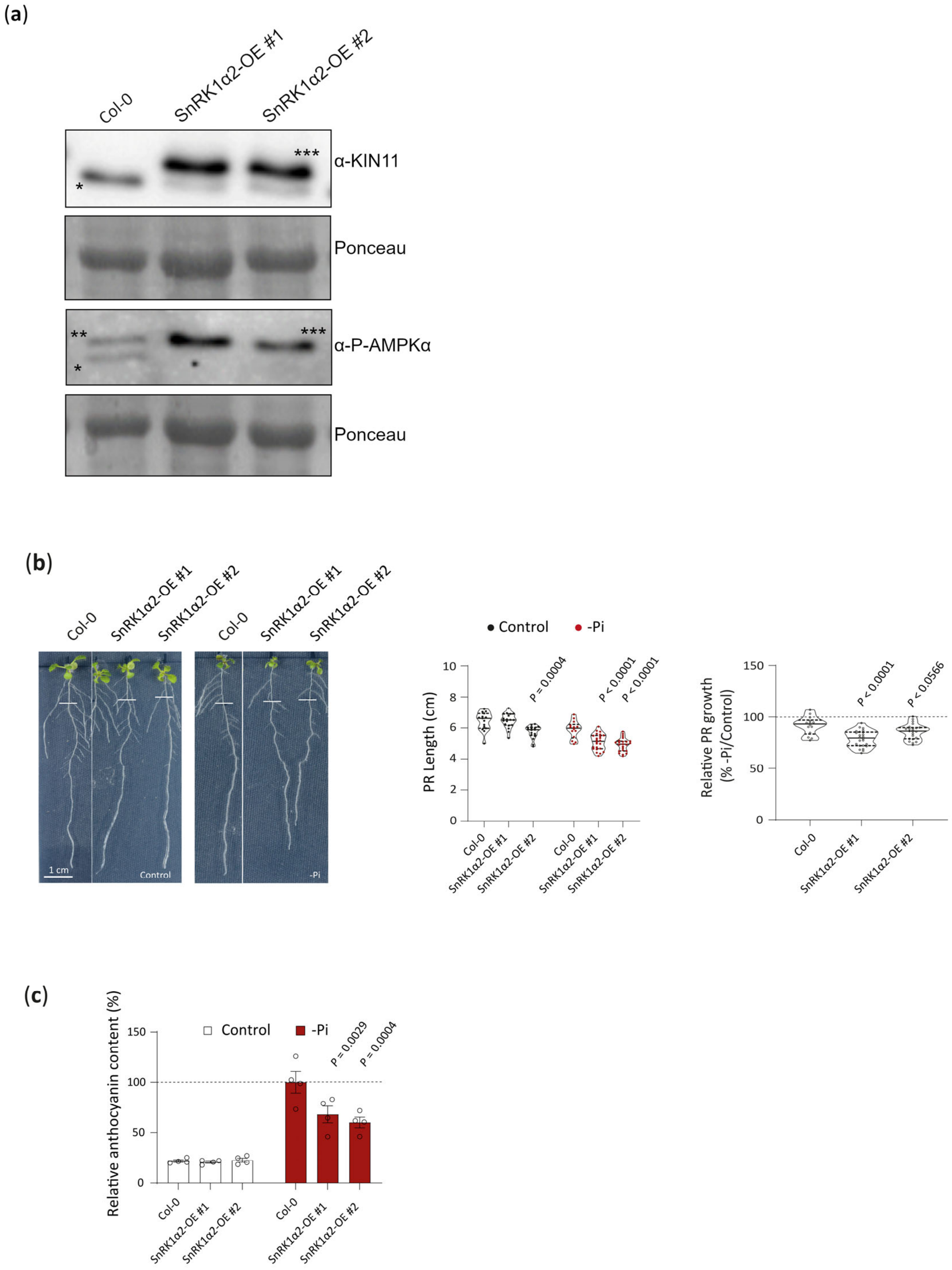


**Figure S4. SnRK1α2 overexpression enhances the inhibition of root growth and attenuates anthocyanin accumulation during phosphate starvation.** (a) Representative immunoblot analysis of KIN11 protein abundance and SnRK1 activation-loop phosphorylation in Col-0 and two independent SnRK1α2-OE lines. Membranes were probed with antibodies against KIN11 and phospho-AMPKα Thr172, which recognises the conserved activation-loop phosphorylation sites of SnRK1α1/KIN10 (Thr175) and SnRK1α2/KIN11 (Thr176). Asterisks indicate the endogenous KIN11 (*), endogenous KIN10 (**), and overexpressed KIN11 (***) signals. Ponceau staining was used as a loading control. (b) Primary root growth of Col-0 and two independent SnRK1α2-OE lines under Pi-sufficient (control) and Pi-deficient (-Pi) conditions. Left, representative images of seedlings after 6 days of vertical growth following transfer to control or -Pi medium. Same images were cropped for showing wild-type (Col-0) and SnRK1α2-OE lines contiguously. Scale bar, 1 cm. Middle, quantification of PR length. n = 14-24 seedlings per genotype and condition. Right, quantification of relative PR growth under -Pi conditions, expressed relative to the corresponding control condition for each genotype (set to 100%). Violin plots show the distribution of individual measurements, with each point representing an individual seedling. *P*-values denote statistically significant differences from wild-type (Col-0) within the same treatment. Two-way (left graph) or one-way (right graph) ANOVA followed by Dunnett's multiple comparisons test. (c) Relative anthocyanin content in Col-0 and two independent SnRK1α2-OE lines grown under control or -Pi conditions for 6 days following transfer to control or -Pi medium. Values are expressed relative to Col-0 under -Pi conditions (set to 100%). Data represent the mean ± SEM. P-values are indicated in the figure. n = 4 biological replicates. *P*-values denote statistically significant differences from wild-type (Col-0) within the same treatment. Two-way ANOVA followed by Dunnett's multiple comparisons test. Col-0 data under control conditions shown in (a) and (b) are the same as those presented in Fig. S2a,b, as the corresponding experiments were performed simultaneously.

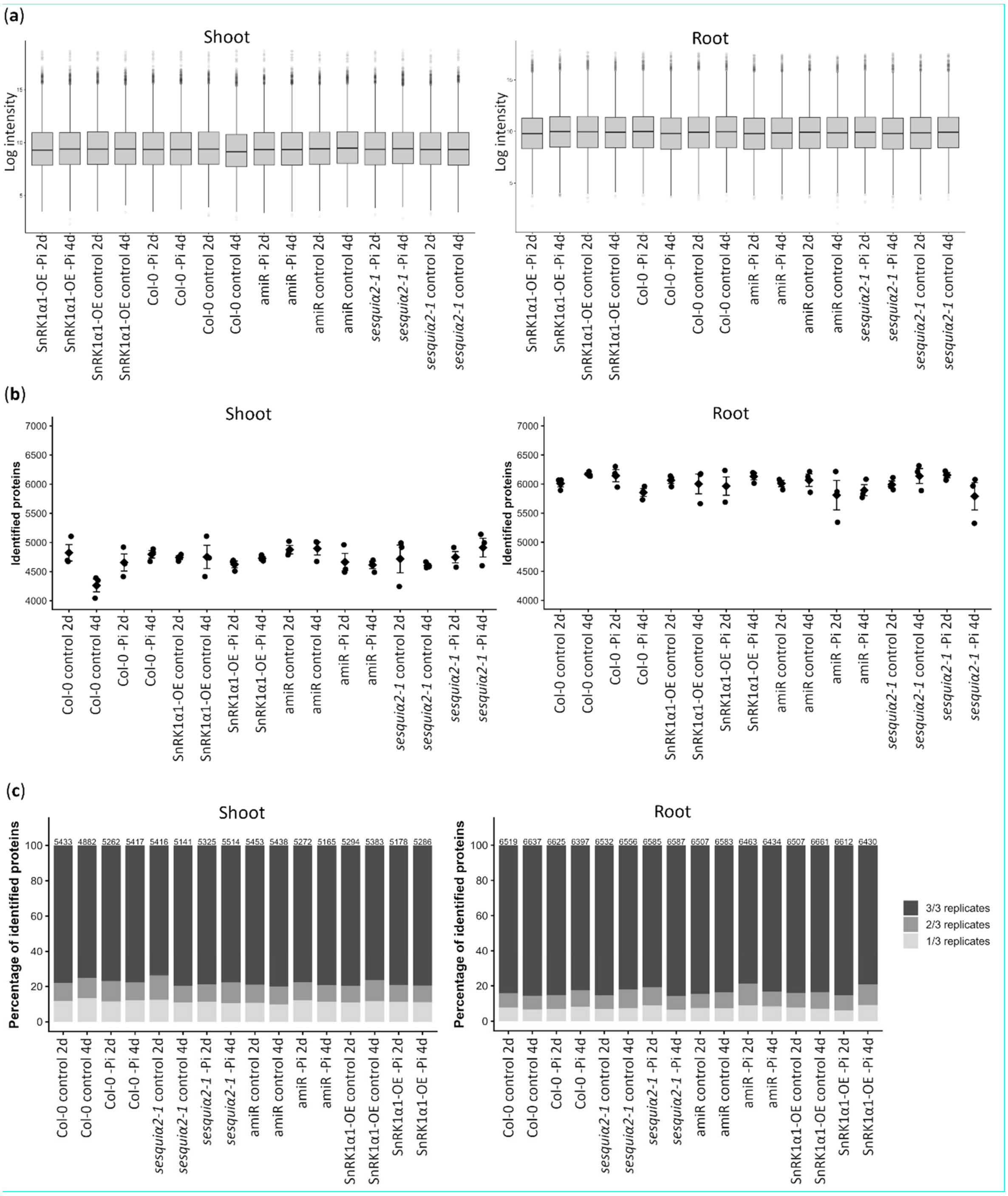


**Figure S5. Quality control of the quantitative proteomic dataset.** (a) Distribution of protein intensities across biological replicates in shoot and root samples. Box plots show the distribution of log-transformed protein intensities for each sample. (b) Number of proteins identified in each biological replicate from shoot and root samples. Data represent the mean ± SEM. Each dot represents one biological replicate (n = 3) and squares represent the mean. (c)

Distribution of proteins identified across biological replicates within each experimental group in shoots and roots. Dark grey, proteins detected in all three biological replicates; light grey, proteins detected in two of three replicates; white, proteins detected in one of three replicates. Numbers above the bars indicate the total number of unique proteins detected across the three biological replicates.

(**a**) Shoot

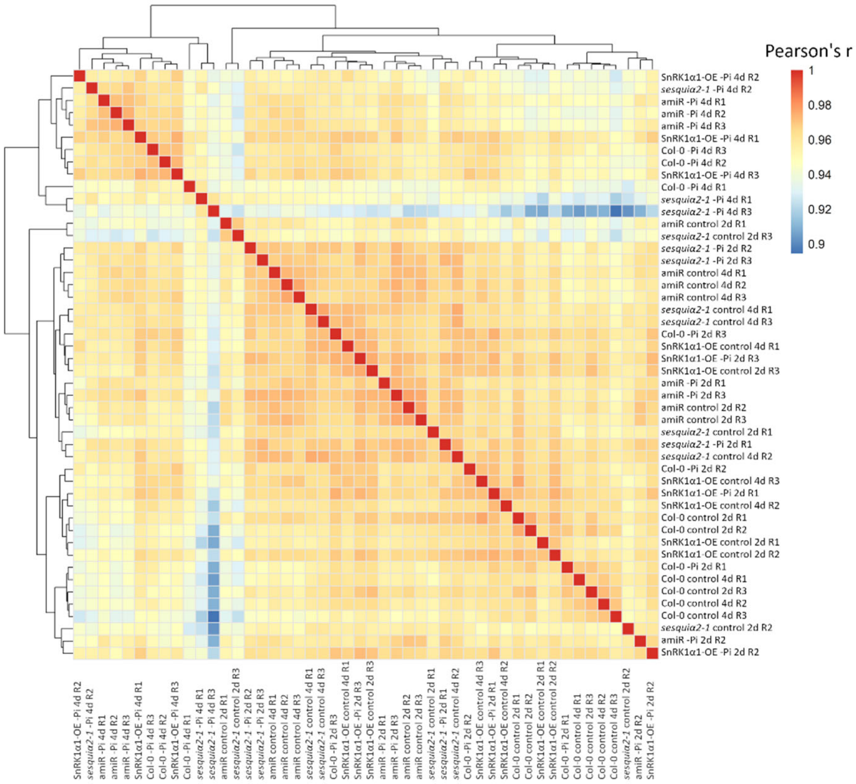


(**b**) Root

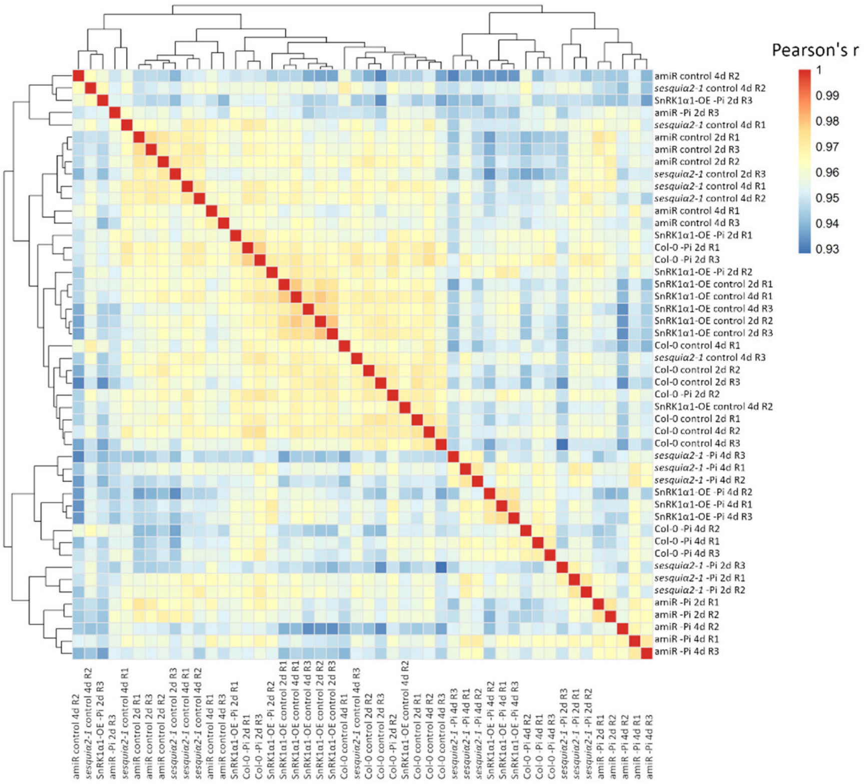

**Figure S6. Reproducibility of the quantitative proteomic dataset.** (a) Pearson correlation matrix showing pairwise correlations between shoot samples. (b) Pearson correlation matrix showing pairwise correlations between root samples. Correlations were calculated based on protein abundance profiles across samples. Colour intensity indicates the Pearson correlation coefficient (*r*), with higher values indicating greater similarity between samples. Hierarchical clustering was performed based on the correlation profiles.

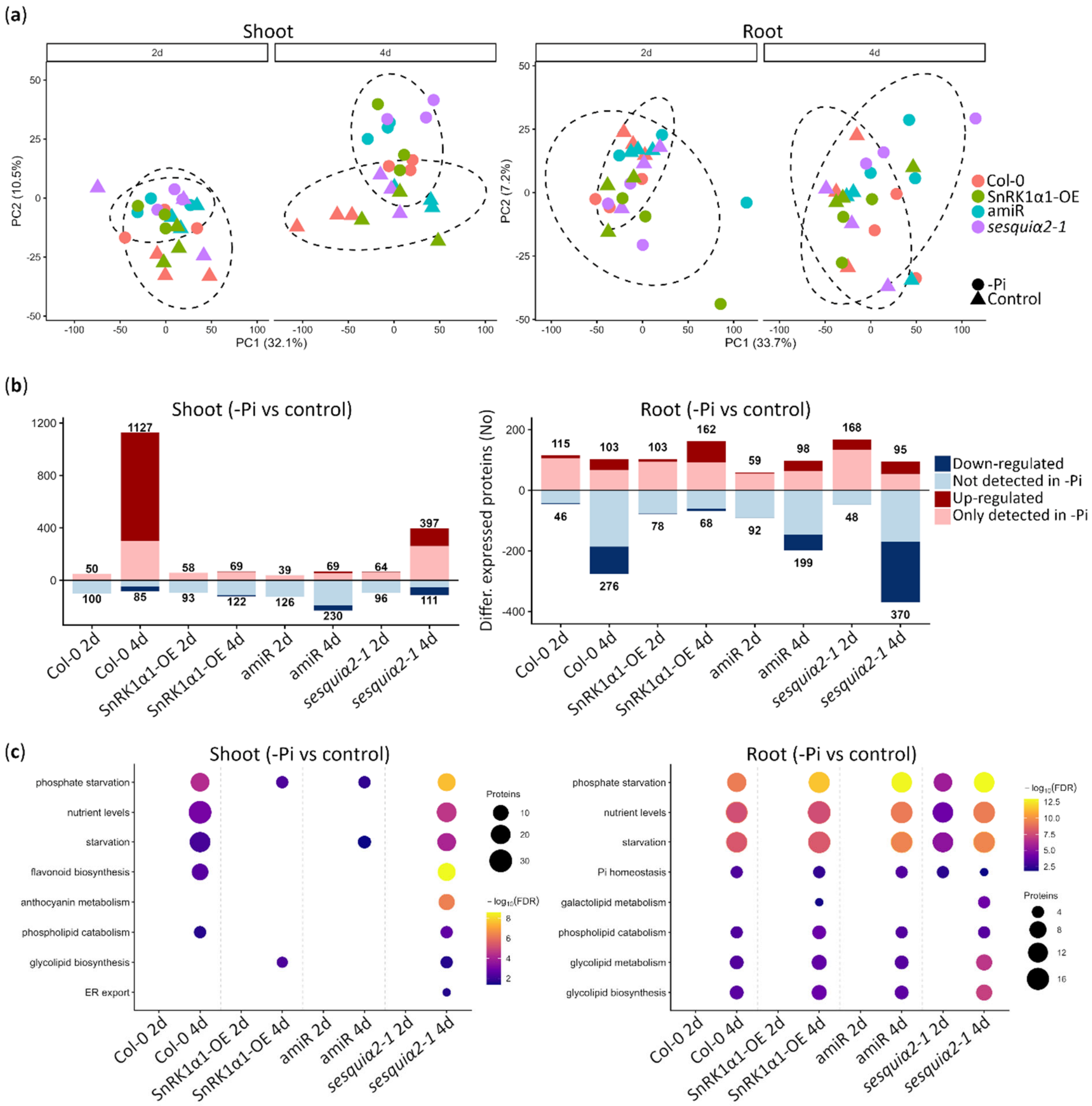


**Figure S7. Global proteomic responses to phosphate starvation.** (a) Principal component analysis (PCA) of shoot and root proteomes from Col-0, SnRK1α1-OE, *amiR-KIN10/11-3* and *sesquiα2-1* seedlings grown under Pi-sufficient (control) or Pi-deficient (-Pi) conditions for 2 or 4 days. Each point represents an individual biological replicate. Dashed ellipses indicate the distribution of samples within the respective groups. (b) Numbers of differentially abundant proteins identified in shoots and roots by comparison of -Pi and control conditions within each genotype at 2 and 4 days. Bars indicate proteins showing decreased abundance under -Pi (blue), and proteins showing increased abundance under -Pi (red), and proteins detected exclusively under one of the two conditions (light blue or right red). (c) Functional enrichment analysis of proteins differentially abundant between -Pi and control conditions for each genotype, tissue and time point. Dot size indicates the number of proteins associated with each enriched category, and colour indicates statistical significance as -log10(FDR).

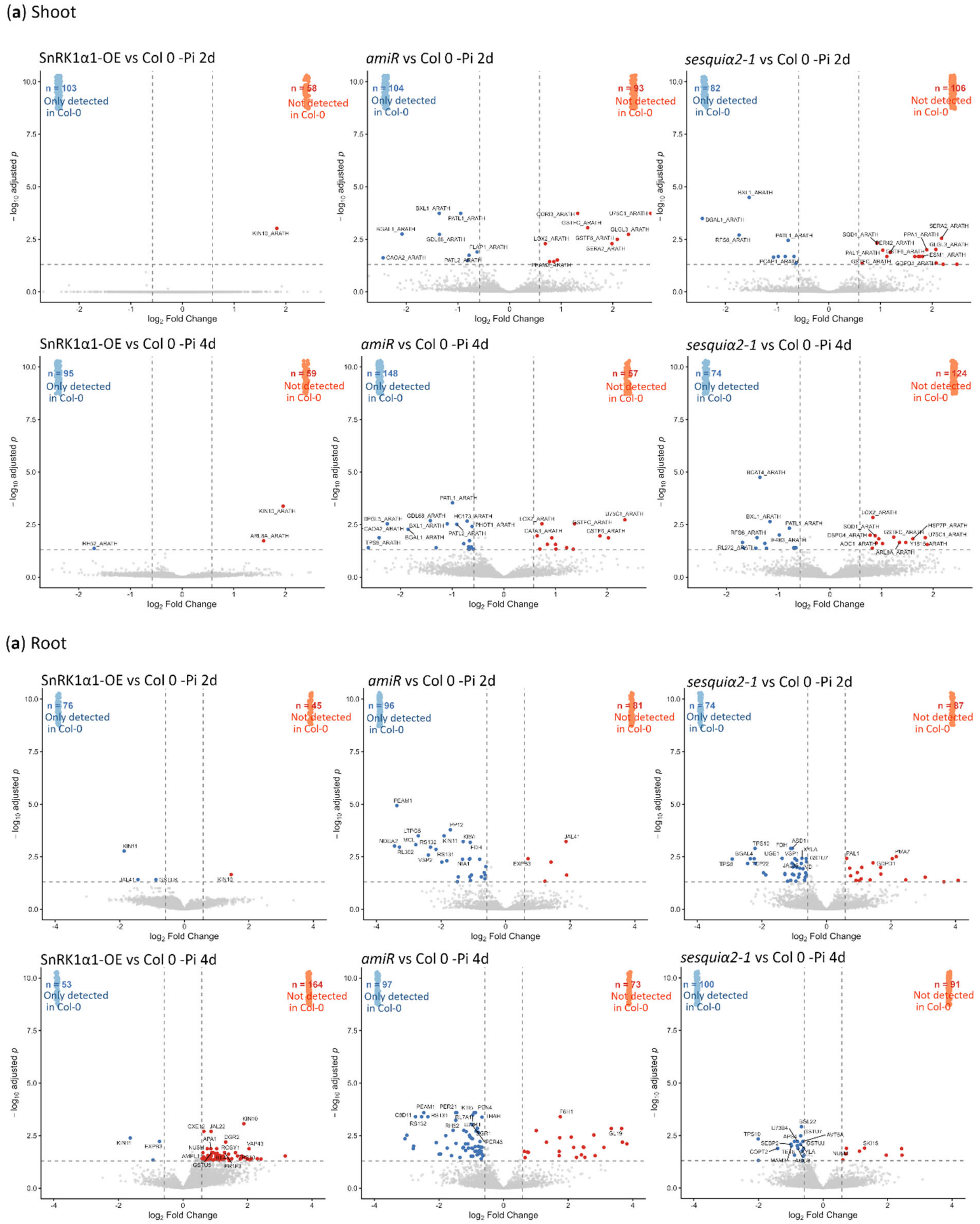


**Figure S8. Genotype-dependent differences in the proteome under phosphate starvation.** (a) Volcano plots showing differential protein abundance between SnRK1-altered genotypes (SnRK1α1-OE, *amiR-KIN10/11-3* and *sesquiα2-1*) and Col-0 under Pi-deficient (-Pi) conditions at 2 and 4 days in shoots. (b) Corresponding analysis in roots.

Each point represents an individual protein. The x-axis shows the $log_2$ fold change and the y-axis the $-log_{10}$(adjusted P-value). Comparisons based on protein detection in only one of three biological replicates in one group versus no detection in the other group (1/3 vs 0/3) were excluded from the analysis. Proteins with an adjusted *P*-value $< 0.05$ and an absolute $log_2$ fold change $> 0.58$ are shown in red (increased abundance) or blue (decreased abundance). Proteins detected in one genotype but not the other are shown separately in lighter shades according to the direction of the difference. The numbers indicate the number of proteins detected exclusively in each genotype. Proteins not meeting these criteria are shown in grey.

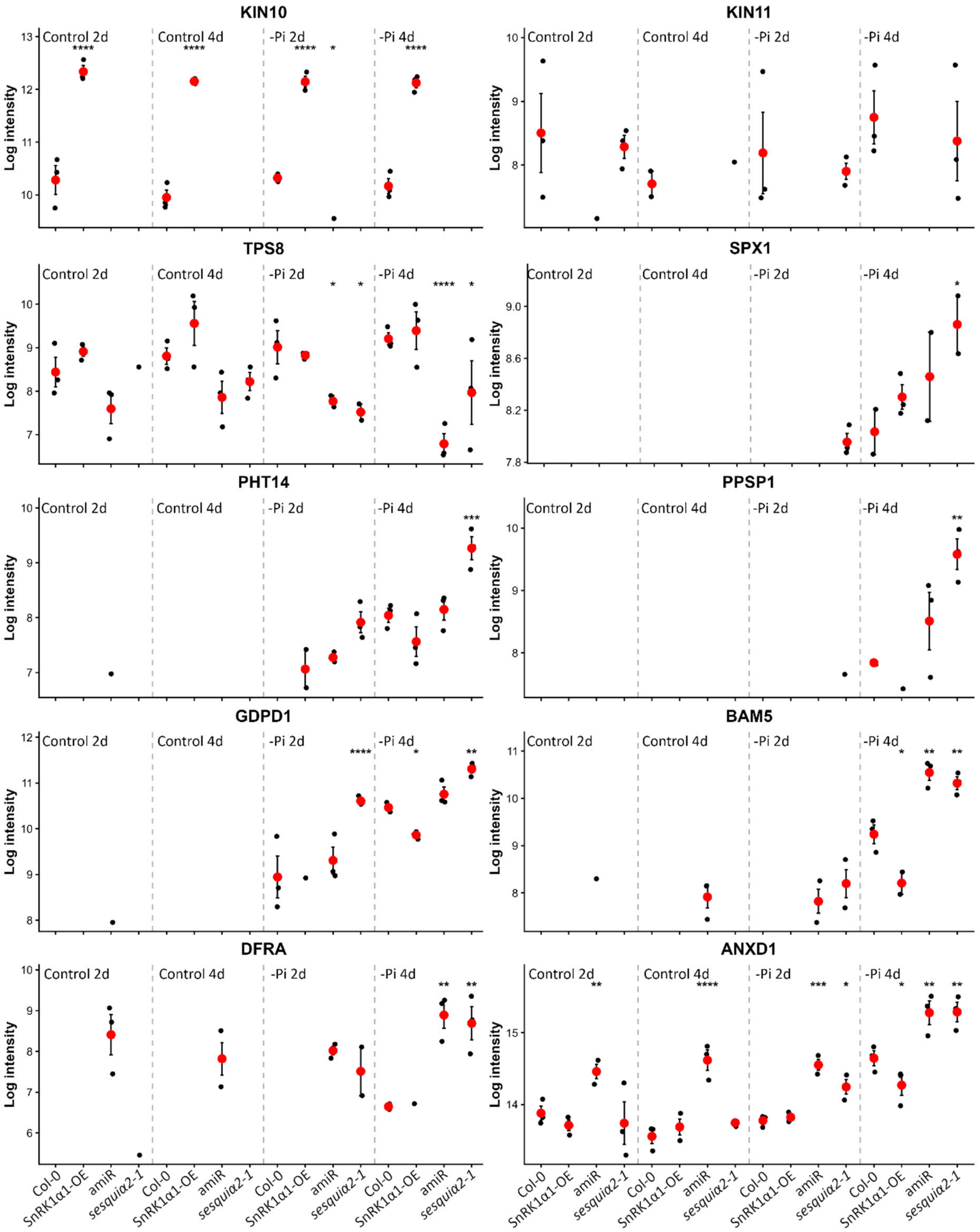


**Figure S9. Shoot abundance of selected proteins associated with SnRK1 signalling and phosphate-starvation responses.** Protein abundance profiles of selected proteins associated with SnRK1 signalling, carbohydrate and phosphate-starvation responses in shoots of Col-0, SnRK1α1-OE, *amiR-KIN10/11-3* and *sesquiα2-1* seedlings grown under Pi-sufficient (control) or Pi-deficient (-Pi) conditions for 2 or 4 days. Each point represents an individual biological replicate, and red points indicate the mean ± SEM. Protein abundance is shown as log-transformed intensity values. Asterisks indicate significant differences between each SnRK1-altered genotype and Col-0 under the corresponding condition and time point, based on MSstats *P*-values. *, $P \leq 0.05$; **, $P \leq 0.01$; ***,$P \leq 0.001$; and ****, $P \leq 0.0001$.

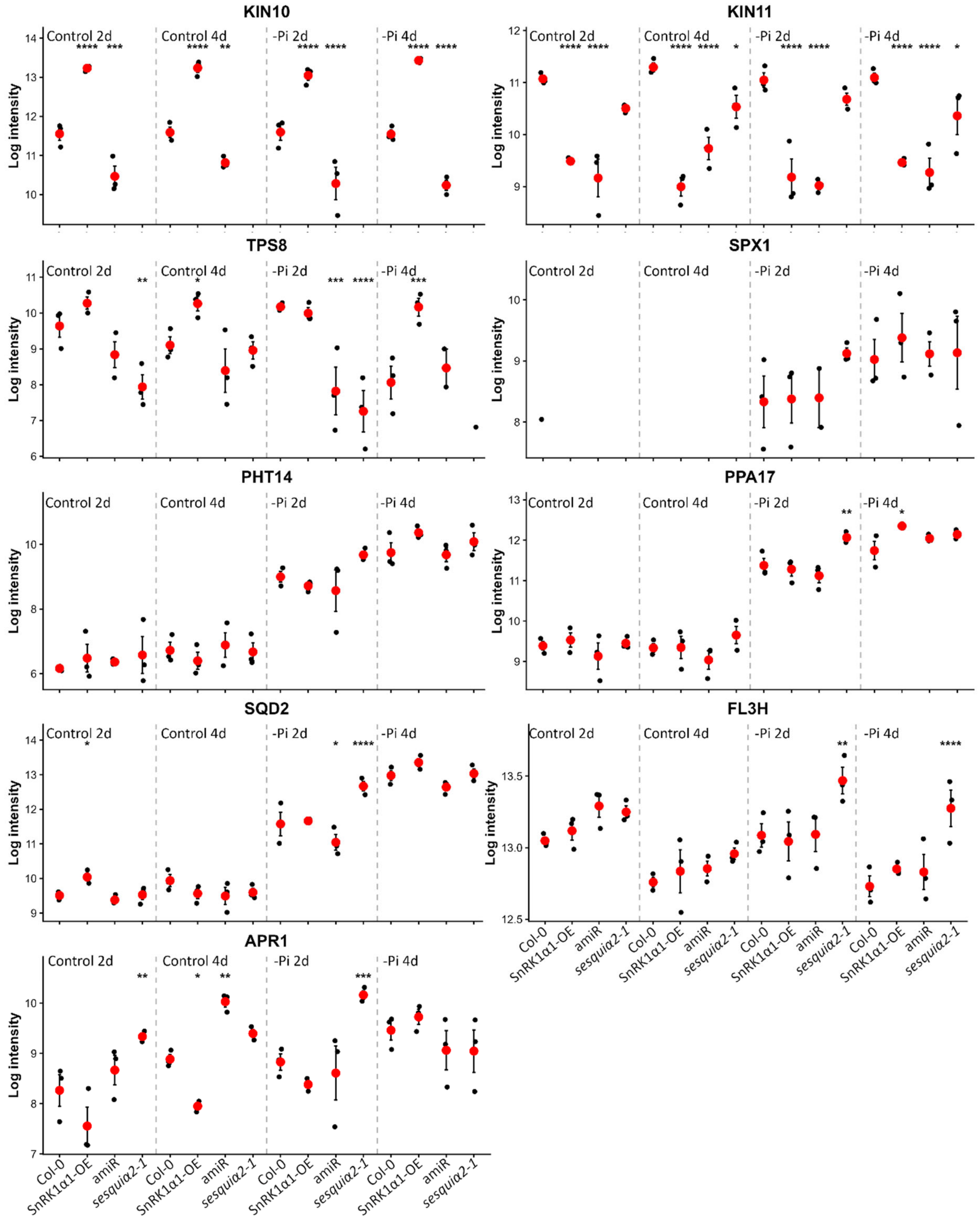


**Figure S10. Root abundance of selected proteins associated with SnRK1 signalling and phosphate-starvation responses.** Protein abundance profiles of selected proteins associated with SnRK1 signalling and phosphate-starvation responses in roots of Col-0, SnRK1α1-OE, *amiR-KIN10/11-3* and *sesquiα2-1* seedlings grown under Pi-sufficient (control) or Pi-deficient (-Pi) conditions for 2 or 4 days. Each point represents an individual biological replicate, and red points indicate the mean ± SEM. Protein abundance is shown as log-transformed intensity values. Asterisks indicate significant differences between each SnRK1-altered genotype and Col-0 under the corresponding condition and time point, based on MSstats *P*-values. *, $P \leq 0.05$; **, $P \leq 0.01$; ***,$P \leq 0.001$; and ****, $P \leq 0.0001$.

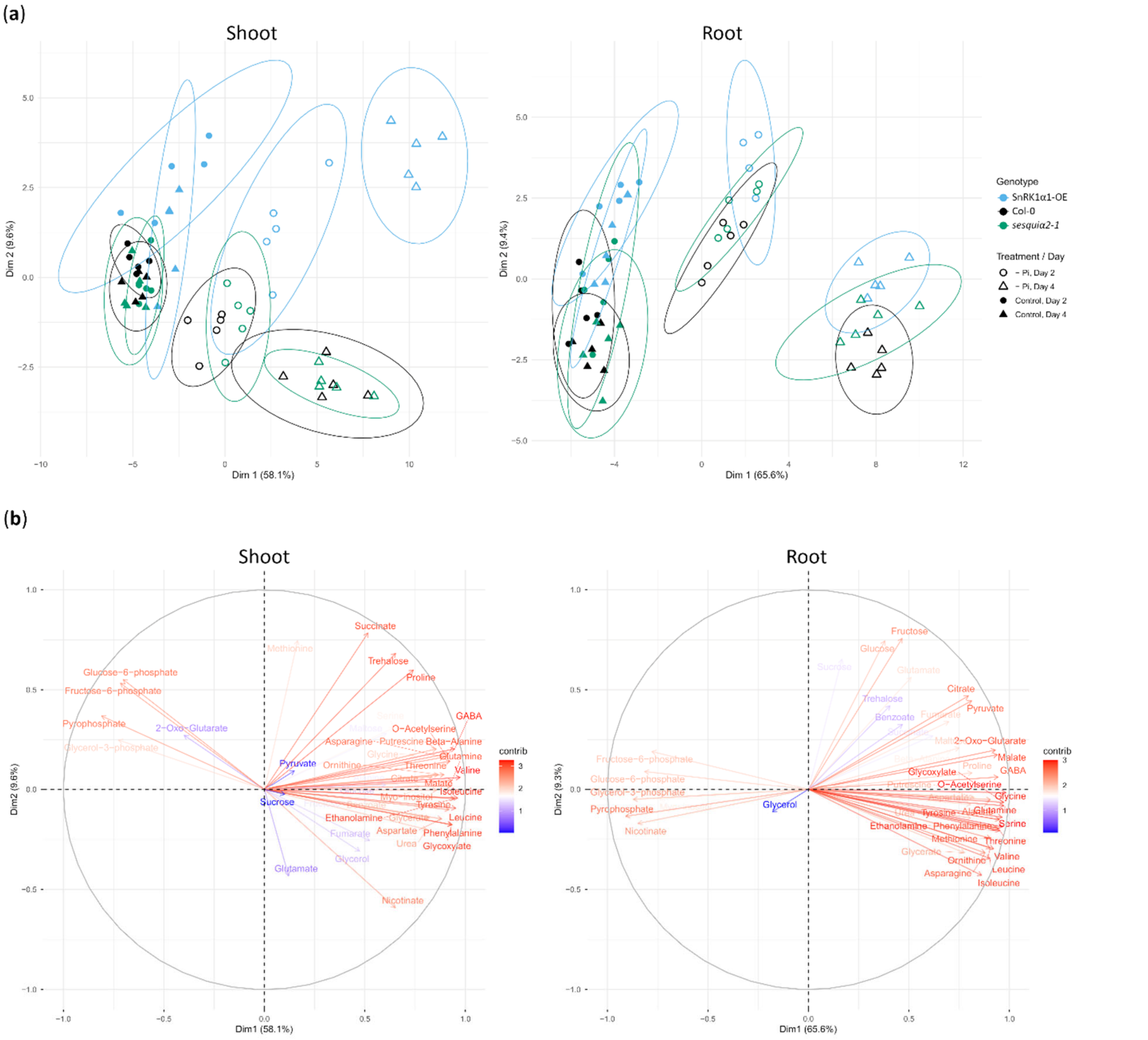


**Figure S11. Principal component analysis of metabolite profiles under phosphate-sufficient and phosphate-deficient conditions**. (a) Principal component analysis (PCA) of metabolite profiles from shoots and roots of Col-0, SnRK1α1-OE and *sesquiα2-1* seedlings grown under Pi-sufficient (control) or Pi-deficient (-Pi) conditions for 2 or 4 days. Each point represents an individual biological replicate. Different colours indicate genotype, and point shapes indicate treatment and sampling time. (b) PCA variable plots showing the contribution and direction of individual metabolites to the first two principal components in shoots and roots. Metabolites are represented by arrows, blue and red colours indicating their contribution to sample separation in the corresponding PCA. The percentages on the axes indicate the proportion of variance explained by each principal component.

**Table S1.** List of the *A. thaliana* lines used in this work

| ***Arabidopsis thaliana* line** | **Genetic background** | **Source** |
|---|---|---|
| *snrk1α1-3* | *GABI_579_E09* | Belda-Palazón *et al.*, 2020 |
| *snrk1α2-1* | *WiscDsLox320B03* | Belda-Palazón *et al.*, 2020 |
| *sesquiα2-1* [*snRK1α1-3 (-/-) α2-1 (+/-)*] | *GABI_579_E09/WiscDsLox320B03* | Belda-Palazón *et al.*, 2020 |
| SnRK1α1-OE | *Col-0* | Jossier *et al.*, 2009 |
| *p35S::ratACC-GFP-HA-HA* | *Col-0* | Muralidhara *et al.*, 2021 |
| *phr1-1* | C663T (Q192*) | Rubio *et al.*, 2001 |
| *ASP-SPARK* | *Col-0* | Safi *et al.*, 2023 |
| *amiR-KIN10/11-1* | *Col-0* | This work |
| *amiR-KIN10/11-2* | *Col-0* | This work |
| *amiR-KIN10/11-3* | *Col-0* | This work |
| SnRK1α2-OE #1 | *Col-0* | This work |
| SnRK1α2-OE #2 | *Col-0* | This work |

**Table S2.** List of primers used in this work.

| **Name** | **Sequence** | **Application** |
|---|---|---|
| amiRKIN10/11-1_1 | gaTGATAGCAACCTTATGCGCTGtctctcttttgtattcc | Generation of an artificial miRNA targeting KIN10/11 in a HBT95-pri319 backbone |
| amiRKIN10/11-1_2 | gaCAGCGCATAAGGTTGCTATCAtcaaagagaatcaatga | |
| amiRKIN10/11-1_3 | gaCAACGCATAAGGTAGCTATCTtcacaggtcgtgatatg | |
| amiRKIN10/11-1_4 | gaAGATAGCTACCTTATGCGTTGtctacatatatattcct | |
| KIN10_RT_Fw | TGCATTTTTTGGTTCCGAATTTTC | RT-PCR analysis |
| KIN10_RT_Rv | GTCCGGTTTAGTATTCAGAGG | RT-PCR analysis |
| KIN11_RT_Fw | GACTTAAAAAAACCTGACTC | RT-PCR analysis |
| KIN11_RT_Rv | ACCCGGCAAAGTAAACACTG | RT-PCR analysis |
| EIF4_RT_Fw | GGCAGTCTCTTCGTGCTGAC | RT-PCR analysis |
| EIF4_RT_Rv | CATGTCTCCGTGAGTTGCAG | RT-PCR analysis |
| SEN5_qPCR_Fw | GCGAAACTCTCTCCGACTTC | RT-qPCR analysis |
| SEN5_qPCR_Rv | CCACAGAACAACCTTTGACG | RT-qPCR analysis |
| DIN1_qPCR_Fw | CAGAGTCGGATCAGGAATGG | RT-qPCR analysis |
| DIN1_qPCR_Rv | ATTTGACCGCTCTCACAACC | RT-qPCR analysis |
| DRM2_qPCR_Fw | CTTCGACAAGCCTTCTCACC | RT-qPCR analysis |
| DRM2_qPCR_Rv | TCGTCGCTGTATAGCCAATC | RT-qPCR analysis |
| PMH1_qPCR_Fw | TCAACGAAATTATGTCATCAGAGG | RT-qPCR analysis |
| PMH1_qPCR_Rv | TTTCAGAGCTCAACATCTTGTCA | RT-qPCR analysis |
| ACP5_qPCR_Fw | CCAAACTTCGAACAATCTTTCTC | RT-qPCR analysis |
| ACP5_qPCR_Rv | GTTTCCCAAAACACTGTACCACT | RT-qPCR analysis |
| SQD1_qPCR_Fw | CATCCTCTAAACCAAAGCGTGT | RT-qPCR analysis |
| SQD1_qPCR_Rv | AGTAGCCCAACCGCAATAAC | RT-qPCR analysis |
| KINβ1_qPCR_Fw | TTATTCGCTCCTCAGGTTCC | RT-qPCR analysis |
| KINβ1_qPCR_Rv | GGTAGGGATTCCTTGCTCTG | RT-qPCR analysis |
| PDF2_qPCR_Fw | TCAACATCTGGGTCTTCACTTAGC | RT-qPCR analysis |
| PDF2_qPCR_Rv | GATGCAATCTCTCATTCCGATAGTC | RT-qPCR analysis |

**Table S3.** List of antibodies used in this work.

| Name | Source | Animal | Dilution |
|---|---|---|---|
| Anti- KIN10 | Agrisera AS10 919 | Rabbit | 1:1000 |
| Anti- KIN11 | Agrisera AS21 4582 | Rabbit | 1:1000 |
| Anti-Phospho-AMPKalpha (Thr172) | Cell Signaling #2531S | Rabbit | 1:1000 |
| Anti Phospho-Acetyl-CoA Carboxylase (Ser79) | Cell Signaling #1673661S | Rabbit | 1:1000 |
| Anti-β Tubulin | Santa Cruz sc-166729 | Mouse | 1:2000 |
| Anti-GFP | Rockland 600-401-215 | Rabbit | 1:5000 |
| Anti-rabbit IgG-HRP | Jacksons #111035144 | Goat | 1:20000 |
| Anti-mouse IgG-HRP | Jacksons #115035146 | Goat | 1:20000 |

**Table S4. Protein abundance measurements used for quantitative proteomic analysis**. Protein abundance data for individual biological replicates from shoots (a) and roots (b) of Col-0, SnRK1α1-OE, *amiR-KIN10/11-3* and *sesquiα2-1* seedlings grown under Pi-sufficient or Pi-deficient conditions for 2 or 4 days. The table also reports the number of measured features, missing values and imputed features for each protein and experimental group.

**Table S5. Differential protein abundance.** Differential protein abundance results generated using MSstatsShiny for shoots (a) and roots (b) across the experimental comparisons. Comparisons based on protein detection in only one of three biological replicates in one group versus no detection in the other group (1/3 vs 0/3) were excluded from the analysis.

**Table S6. Functional enrichment analysis of differentially abundant proteins.** Gene Ontology (GO) enrichment results for proteins showing differential abundance across the indicated comparisons, tissues and directions of change. *P*-values were adjusted for multiple testing using the Benjamini-Hochberg method.

**Table S7. Primary metabolite abundance data.** Primary metabolite abundance data for individual biological replicates from shoots and roots of Col-0, SnRK1α1-OE and *sesquiα2-1* seedlings grown under Pi-sufficient or Pi-deficient conditions for 2 or 4 days. The table reports the relative abundance of individual metabolites for each biological replicate together with the corresponding metabolite annotation and experimental condition. Quality-control samples are indicated as QC.

**Table S8. Statistical analysis of metabolite abundance differences shown in Figure 4.** Statistical results for the comparison of primary metabolite abundance between Col-0 and SnRK1-altered genotypes in shoots (a) and roots (b) under Pi-sufficient (control) and Pi-deficient (-Pi) conditions after 2 or 4 days of treatment. *P*-values were calculated using Welch's t-test and adjusted using the Benjamini-Hochberg method.
